# Stop-Flow Lithography Using Hydrodynamic Flow Resistance Regulation and Hydraulic Compliance Minimization for Size-Tunable Anisotropic Multilayered Microparticles Fabrication

**DOI:** 10.64898/2026.11.01.763425

**Authors:** Yiying Zou, Muhammad Usman Akhtar, Ghulam Destgeer

## Abstract

Stop-flow lithography (SFL) enables the fabrication of microparticles with designed geometries and material compositions, but its performance relies on rapid flow stopping, which remains particularly challenging in multi-inlet systems with high hydrodynamic flow resistance and hydraulic compliance. Here, we develop a pneumatic-controlled SFL platform with a framework based on flow resistance regulation and hydraulic compliance minimization for fabricating size-tunable anisotropic multilayered microparticles. Our analysis indicates that flow-stopping behavior is governed by the release of extra fluid volume stored in the system due to hydraulic compliance, which arises from the deformation of microfluidic components and the compressibility of the fluid within them. Through numerical modeling and analysis, we identify deformation in the PDMS inlet port region as the dominant contributor to the stored extra fluid volume. By regulating the flow resistance of the tubing relative to that of the outlet channel, the stored extra fluid volume can be preferentially released forward through the outlet or backward through the inlet tubing, enabling rapid flow stopping across configurations with 500, 250, and 100 μm outlets. Furthermore, by supporting the PDMS membrane beneath the inlet region with a rigid glass substrate and reducing its dimensions, the hydraulic compliance and consequently the stored extra fluid volume are significantly reduced. With these strategies, microparticles with well-defined boundaries can be fabricated at short, flow-stopping, delay times, and the flow profiles obtained in multi-inlet systems can be preserved during flow stopping and transferred into particle patterns.

## 1. Introduction

Microfluidic technology derives its unique advantages from the ability to precisely manipulate fluid flows at the microscale. Substantial progress has been achieved in the control of steady flows under constant flow rate conditions, enabling applications such as concentration gradient generation^1^, uniform droplet formation^2,3^ and coalescence^4^, as well as the sorting of droplets^5^, microparticles^6–8^ and cells^9^. Beyond steady flow control, an increasing number of emerging applications require precise control of dynamic flows^10^. Such dynamic flow controls are particularly important in applications including pulsatile shear flow stimulation for endothelial cell mechanotransduction^11^, multi-material switching in micro-fabrication^12–18^ and 3D printing^19–21^.

Stop-flow lithography (SFL)^12–16^ is a representative application in which reliable dynamic stop-flow control is critical. As a fundamental step of the dynamic stop-flow control, the flow-stopping process significantly affects the quality of fabricated microparticles and is more complex than simply removing the driving source. Ideally, the flow should immediately stop once the driving source is removed. In practice, however, flow stopping is usually delayed because a certain amount of extra fluid volume is stored in the microfluidic circuit when the system is pressurized. This pressure-dependent fluid volume change Δ*V* is associated with the hydraulic compliance *C*, defined as 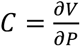 Upon removal of the driving source, the stored fluid volume change will be released and act as a transient driving force, generating residual flow until it is fully discharged, after which the flow eventually stops.

The stored volume change can arise from two major effects. The first is the deformation of the microfluidic components, particularly the elastic PDMS channels^12,22^. Under pressurized condition, the walls of the PDMS channel deform, thereby expanding the fluid volume stored within the circuit. This stored volume change will be released when the channel walls recover upon depressurized condition. Although this effect is inevitable in elastic PDMS channel, the ease of fabrication and high oxygen permeability still make PDMS channel the most used device for flow lithography. Oxygen permeability is important for preventing channel clogging during particle fabrication and eliminates the need for sheath flow in flow lithography. The second effect is the compressibility of the fluid dead volume inside the microfluidic components, which gives rise to the so-called “bottleneck effect”^23^. This effect usually occurs when the syringe pump is used as the driving source. As the syringe plunger advances, pressure gradually builds up to overcome the flow resistance of the microchannel. During this process, part of the plunger displacement is accommodated by compression of the fluid rather than being immediately delivered through the microchannel, resulting in an extra compressed fluid volume stored within the circuit, which will be released upon depressurized condition. The bottleneck effect usually becomes more pronounced with a larger fluid dead volume inside the syringe or higher microchannel flow resistance. In the syringe-pump cases investigated in ref. [23], the fluid volume change caused by the bottleneck effect contributed more significantly to delayed flow response than that arising from the flexibility of microfluidic components.

To mitigate these issues and achieve rapid flow stopping, syringe-pump systems commonly employ valves to suppress or release that extra fluid volume^13–15,24–27^. Nevertheless, even equipped with valves, syringe-pump systems still underperform in high-flow-resistance microchannels^28^. In contrast, pneumatic-pump systems offer greater potential for rapid flow stopping in high-flow-resistance microchannels because they are not affected by the bottleneck effect^23^ and thus usually exhibit shorter response times^29^. However, these systems suffer from another limitation: hydrostatic pressure imbalance arising from height differences between the liquid levels in reservoirs and the channel outlet can induce unintended hydrostatically driven flow even when the driving source is removed^28,30^. Over extended operation, the liquid levels change over time, requiring the continuous adjustment of the reservoir heights relative to the channel outlet.

Moreover, rapid stopping of multiple co-flowing streams within microchannels is particularly important for the fabrication of multi-material microparticles. Excessively long flow-stopping delays generally cause serious diffusion-induced material mixing issues and destruction of the established flow profiles. Although various methods have enabled versatile flow-profile sculpting, rapidly stopping the multiple co-flowing streams in high-flow-resistance microchannels and faithfully preserving the established flow profiles in the final particle architectures remains a challenge in SFL^27^.

In this work, we present a pneumatic-controlled stop-flow lithography (SFL) platform for the fabrication of size-tunable anisotropic multilayered microparticles (**Fig. 1**). To mitigate the hydrostatic pressure imbalance issue inherent in pneumatic-controlled microfluidic systems, we designed five channel configurations by systematically regulating the flow resistance of the tubing relative to that of the outlet channel, thereby limiting unintended hydrostatically driven flow to an acceptable level (**Table 1**, configs. 1–5). Guided by the hydraulic–electric analogy, we investigate the generation and release of extra fluid volume stored in the microfluidic circuit by using numerical modeling and analysis in both single-inlet configurations and multi-inlet configurations. In single-inlet configurations, our simulations indicate that the deformation of the inlet port is the dominant contribution to the stored extra fluid volume and further investigate its deformation mechanism and influencing factors, including the applied pressure, PDMS stiffness, and channel configuration in our system. Depending on the relative flow resistances of the tubing and outlet in different channel configurations, the stored extra fluid volume can be rapidly released in either the forward or backward direction, enabling the fabrication of single-material microparticles with tunable sizes and well-defined boundaries. Extending the analysis to multi-inlet configurations, we further explore strategies to reduce the deformation of the inlet port section by supporting the PDMS membrane with a rigid glass substrate and reducing the dimensions of the inlet port section. These results provide practical design rules for minimizing flow-stopping delay in SFL and patterned microparticles were experimentally fabricated with the faithful preservation of the profiles established in multi-inlet configurations.

**Figure 1.**
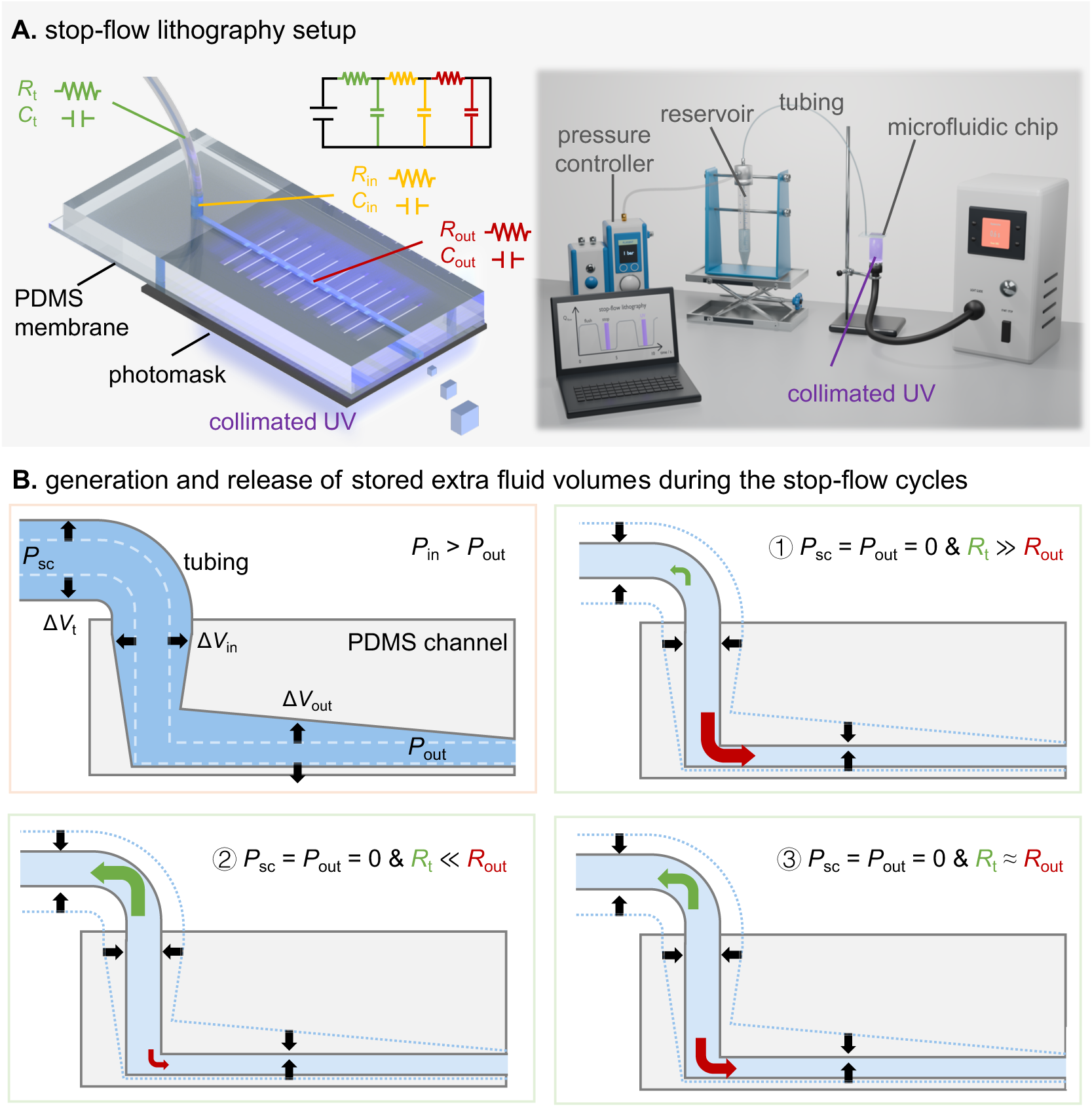
Pneumatic-controlled stop-flow lithography (SFL) platform for fabricating microparticles with tunable sizes. (A) Left: Illustration of a single-inlet microfluidic chip based on the hydraulic–electric analogy, in which hydrodynamic flow resistance *R* (tubing resistance *R*t, inlet port resistance *R*in, and outlet resistance *R*out) and hydraulic compliance *C* (tubing compliance *C*t, inlet port compliance *C*in, and outlet compliance *C*out) are analogous to the electrical resistance and capacitance, respectively. Right: SFL setup consisting of a pneumatic controller, reservoir, tubing, microfluidic chip with photomask underneath, and UV light unit (back row, from left to right), with a laptop (front row) sending synchronized signals to the pneumatic pump and UV light unit. A 3D-printed adapter used to position the chip and prevent UV leakage is omitted for clarity. (B) Flow stopping governed by the generation (light pink frame) and release (light green frame) of extra fluid volumes stored within the microfluidic circuit. Under pressurized condition (*P*sc > *P*out), extra fluid volumes are generated (volume changes Δ*V*) and stored within the microfluidic circuit due to the corresponding hydraulic compliances *C* in (A). The white dashed lines indicate the original channel boundaries before pressurization, and the medium-blue region represents the compressed state of the fluid. Under depressurized condition (*P*sc = *P*out = 0), the stored Δ*V* is released along different flow paths depending on the relative flow resistances of tubing *R*t and outlet *R*out: primarily ① through the outlet when *R*t ≫ *R*out, ② through the tubing when *R*t ≪ *R*out, and ③ through both the tubing and outlet when *R*t ≈ *R*out. The blue dotted lines indicate the expanded channel boundaries at pressurized condition, and the light-blue region represents the decompressed state of the fluid. Green and red arrows indicate fluid release direction toward the tubing and outlet, respectively.

**Table 1.** Channel configurations.

| config. | outlet |  | tubing |  | hydrostatic velocity (µm/s) | *R*<sub>t</sub> *:* *R*<sub>out</sub> | volume release |
|---|---|---|---|---|---|---|---|
|  | *W* × *H* × *L*<br>(mm ×mm × mm) | *R*<sub>out</sub><br>(Pa.s/m<sup>3</sup>) | *I.D.* × *L*<br>(mm × mm) | *R*<sub>t</sub><br>(Pa.s/m<sup>3</sup>) |  |  |  |
| 1 | 0.5*0.5*35 | 1.20 × 10<sup>11</sup> | 0.25*600 | 4.72 × 10<sup>13</sup> | 8.19 | 394:1 | forward |
| 2 | 0.25*0.25*35 | 1.92 × 10<sup>12</sup> | 0.18*600 | 1.76 × 10<sup>14</sup> | 8.73 | 92:1 | forward |
| 3 | 0.1*0.1*35 | 7.50 × 10<sup>13</sup> | 0.13*600 | 6.46 × 10<sup>14</sup> | 13.45 | 8.6:1 | forward |
| 4 |  |  | 0.25*600 | 4.72 × 10<sup>13</sup> | 79.32 | 1:1.6 | forward & backward |
| 5 |  |  | 0.50*600 | 2.95 × 10<sup>12</sup> | 124.36 | 1:25.4 | backward |

\* Estimated with a working fluid with density *ρ* = 987 kg/m<sup>3</sup> and viscosity *μ* = 7.55 mPa·s, using an online microfluidic calculator provided by Elveflow (https://www.elveflow.com/microfluidic-calculator/). Hydrostatic velocity is estimated with 1 cm height difference between the liquid levels in reservoirs and the channel outlet (*P*<sub>h</sub> = *ρgh* = 0.97 mbar, *g* = 9.8 m/s²) in single-inlet configurations.

## 2. Results and Discussion

The objective of this study is to understand and optimize the flow-stopping process in SFL and thus facilitate the fabrication of high-quality microparticles. As discussed in the introduction, the delay of flow stopping is mainly caused by the release of the extra fluid volume stored in the microfluidic circuit. Here, we adopt the hydraulic–electric analogy^31^ to understand the generation and release of the stored extra fluid volume in our microfluidic system (**Fig. 1**): in pressurized condition, the pressure difference across the flow channel acts like a voltage source in the electric circuit, driving fluid flow while storing extra fluid volume (volume change Δ*V*) through system hydraulic compliance *C*, analogous to storing the charge through electrical capacitance; in depressurized condition, the Δ*V* is released through different flow paths depending on the hydrodynamic flow resistance *R*, analogous to discharge through electrical resistance. Contributions to the total volume change Δ*V*_total_ were estimated by considering two mechanisms (material deformation and fluid compressibility) across three components (PEEK tubing, PDMS inlet port, and PDMS outlet). These contributions comprise Δ*V*_t, m_, Δ*V*_t, f_, Δ*V*_in, m_, Δ*V*_in, f_, Δ*V*_out, m_, and Δ*V*_out, f_. The material deformation-induced Δ*V* were obtained from COMSOL simulations, while those arising from fluid compressibility were calculated theoretically (see Materials and Methods for details).

We first investigated the generation of Δ*V* in our single-inlet configurations (**Fig. 2A** and **Table 1**, same inlet port geometry with 500, 250, and 100 μm outlets for configs. 1,2, and 3–5, respectively). Under an applied source pressure *P*_sc_ of 2 bar, Δ*V*_total_ of single-inlet configurations increases from 0.125 μL in config. 1 to 0.221 μL in config. 2 and 1.343 μL in config. 3 (**Fig. 2B** and **Table S3**). This pronounced increase is largely associated with the increase of volume change from the PDMS inlet port (Δ*V*_in, m_, red text, medium-gray region), which dominates Δ*V*_total_ among the six contributions. The increase of Δ*V*_in, m_ among configs. 1–3 can be attributed to the increase of the actual pressure distributed in the inlet port *P*_in_. Specifically, under *P*_sc_ of 2 bar, *P*_in_ values were approximately 0.005, 0.022, and 0.208 bar for the configs.1, 2, and 3 respectively, estimated according to the regulated flow resistance ratios between the tubing and outlet (**Table 1**). The higher *P*_in_ then results in substantially higher Δ*V*_in, m_ and thus higher Δ*V*_total_. Configs. 3–5 are designed to have the same 100 μm outlet but employ tubing with different flow resistances, which results in similar *P*_in_ at source pressures *P*_sc_ of 2, 0.34, and 0.215 bar, respectively (**Table S1**).

**Figure 2.**
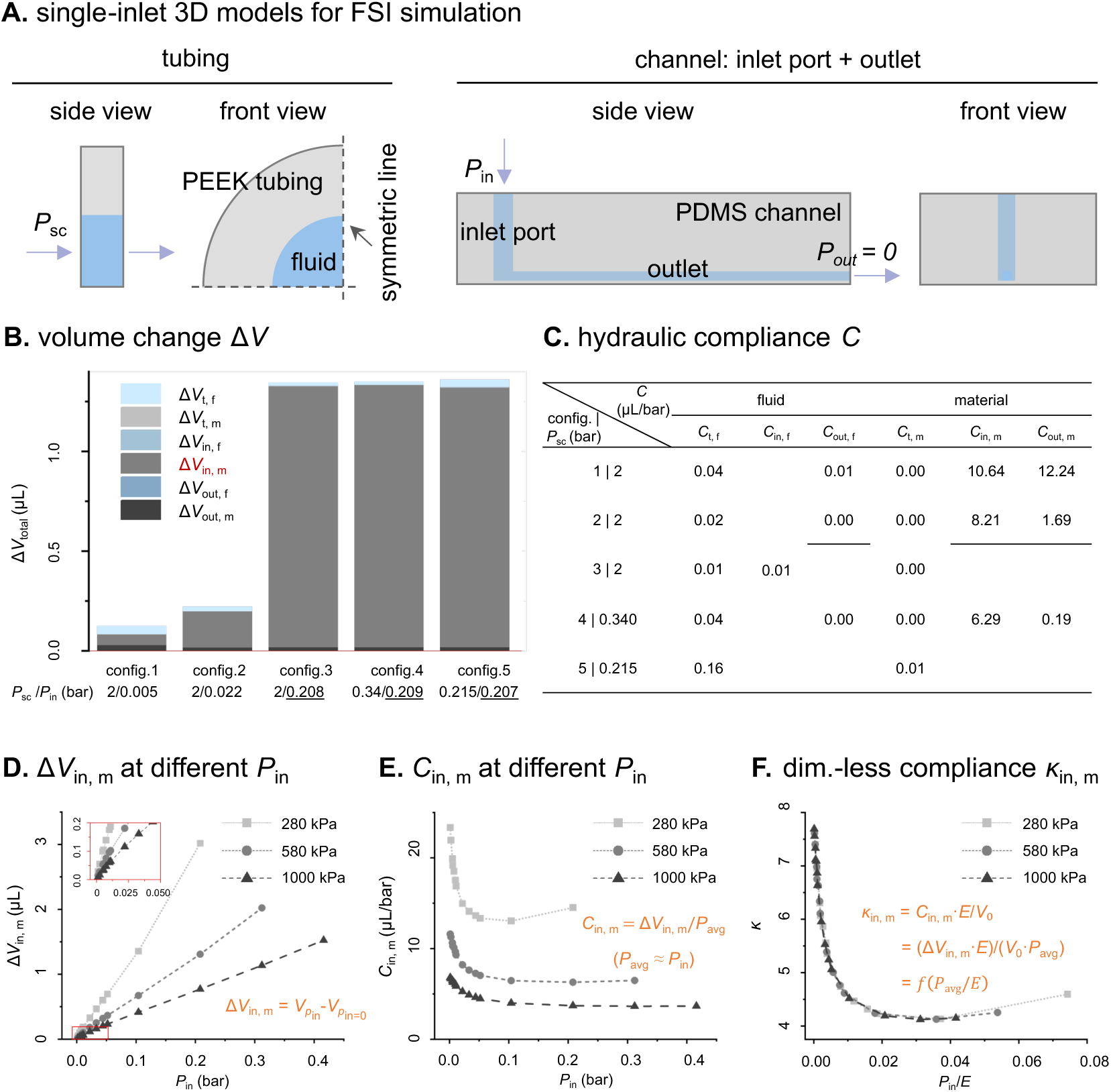
Numerical modeling and analysis of volume changes Δ*V* and hydraulic compliances *C* in single-inlet configurations. (A) 3D models used for fluid–structure interaction (FSI) simulations, including a quarter-symmetry model of the PEEK tubing and a full model of PDMS channel comprising the inlet port and outlet. Corresponding boundary conditions are provided in **Table S2**. (B) Δ*V*total and their six contributions arising from material deformation (m) and fluid compressibility (f) in the PEEK tubing (t), PDMS inlet port (in), and PDMS outlet (out) for configs.1–5. The applied source pressure *P*sc and corresponding pressure distributed in inlet *P*in are indicated below the bars. Δ*V* data is provided in **Table S3**. (C) Secant hydraulic compliances *C* corresponding to the Δ*V* in (B), calculated by dividing the magnitude of each Δ*V* by their corresponding average pressure *P*avg in the respective component (**Tables S1&S3**). (D) Δ*V*in, m as a function of *P*in for three PDMS Young’s moduli (E = 280, 580, and 1000 kPa). The inset zooms in on the low-pressure region. (E) Secant Cin, m as a function of *P*in for three PDMS Young’s moduli. (F) Dimensionless secant compliance κ plotted against normalized inlet pressure *P*in/*E*, showing the approximate collapse of the results for all three Young’s moduli onto a common curve. *P*in values were determined from configs. 1–3 under *P*sc of 0.5, 1, 2, 3, and 4 bar. Since the pressure drop in inlet port is negligible compared to other components, *P*avg was assumed to be equal to *P*in here.

To further understand the underlying physical mechanisms, we also analyzed the fluid compressibility-induced compliance *C*_f_ and material deformation-induced compliance *C*_m_ corresponding to each Δ*V* contribution shown in **Fig. 2B** (**Fig. 2C**). *C*_f_ can be estimated with *C*_f_ = Δ*V*/*P*_avg_ = *V*_0_/*K*_e_, where *V*_0_ is the fluid dead volume and *K*_e_ is the effective bulk modulus of the fluid. Assuming a nearly constant *K*_e_ over the temperature and investigated pressure range in our study, *C*_f_ is therefore governed by the fluid dead volume accommodated by each microfluidic component and is approximately independent of pressure. In contrast, *C*_m_ is more complicated and influenced by structural geometry, elastic modulus, and pressure. C_in, m_ decreases from 10.64 μL/bar in config. 1 to 8.21 μL/bar in config. 2 and 6.29 μL/bar in config. 3 corresponding to the increased *P*_in_. C_out, m_ exhibits a much more pronounced decrease across the configurations than that of C_in, m_. In addition to the effect of increased *P*_in_, this pronounced decrease of C_out, m_ can be attributed to changes in the outlet geometry: decreasing the channel width and height can significantly reduce the channel compliance^23^.

To better understand the pressure-dependent deformation behavior of the inlet port, we numerically investigated the effects of *P*_in_ and PDMS Young’s modulus *E* on Δ*V*_in, m_ and corresponding *C*_in,m_. As shown in **Fig. 2D**, Δ*V*_in, m_ exhibits a nonlinear increase with the increase of *P*_in_ under three Young’s moduli, with softer PDMS exhibiting higher Δ*V*_in, m_ at a given *P*_in_. We further calculated *C*_in,m_ and found that *C*_in,m_ decreases sharply with the increase of *P*_in_ at low pressures, approaches a relatively stable value at intermediate pressures, and exhibits a slight increase at higher pressures (**Fig. 2E**). When *C*_in,m_ is normalized as dimensionless compliance *κ*_in,m_ = *C*_in,m_·*E*/*V*_0_, where *V*_0_ is the initial fluid volume of the inlet port (8.82 μL), and plotted against the normalized inlet pressure *P*_in_/*E*, the curves for all three moduli approximately collapse into a single curve (**Fig. 2F**). As expected from dimensional analysis for a fixed geometry and Poisson’s ratio, this collapse reveals a common scaling relationship between Δ*V*_in, m_ and *P*_in_/*E*, and the persistent shape of the curves suggest a nonlinear pressure-dependent deformation mechanism for the investigated inlet port geometry, potentially contributed by a large PDMS membrane deflection at the bottom of the inlet port, including membrane bending and stretching^32^ (**Fig. S3**). Further controlled simulations with a rigid glass substrate supporting the bottom PDMS membrane show that *C*_in,m_ significantly decreased and became nearly pressure-independent (increasing by about 12% up to ∼0.3 bar) when large membrane deflection is suppressed, indicating an approximately linear relationship between Δ*V*_in, m_ and *P*_in_/*E* under this situation (**Fig. S4**). The slight increase of *C*_in,m_ at the highest pressures (*P*_in_/*E* ≳ 0.04; about 10% for *E* = 280 kPa) also appears in the glass-supported case (Fig. S4), indicating that it originates mainly from the roof and sidewalls of the inlet port rather than from the floor membrane. We attribute it to large-deformation softening of the pressurized PDMS cavity, analogous to the inflation of a balloon: as the cavity expands, the loaded surface grows and the surrounding PDMS thins, so that each pressure increment produces a larger volume change. The magnitude of this softening may depend on the constitutive model used for PDMS (Section 4.4).

The release of Δ*V* governs the flow-stopping process in SFL and consequently affects the quality of the fabricated microparticles. Upon removal of the applied source pressure, Δ*V* stored through the hydraulic compliance is gradually released, sustaining a transient flow until the flow fully stops. During this process, *t*_stopping_ is defined as the time required for the flow to fully stop following pressure removal, whereas *t*_delay_ refers to the preset interval between pressure removal and the onset of UV exposure in SFL process. When *t*_delay_ ≥ *t*_stopping_, microparticles with well-defined boundaries can be fabricated, whereas *t*_delay_ < *t*_stopping_ may result in particles with blurred boundaries or excessive thickness. Single-inlet configurations with 500 and 250 μm outlets allowed short *t*_delay_ of 0.1 and 0.4 s, respectively, for well-defined particle fabrication, whereas the configuration with 100 μm outlet required a prolonged *t*_delay_ of 8 s (**Fig. 3**, configs. 1–3, under *P*_sc_ of 2 bar). The flow-stopping process depends not only on the amount of Δ*V* that needs to be released, but also on its release rate, which is affected by the dimension of the available release flow paths. The 8 s *t*_delay_ required for the 100 μm outlet configuration therefore is consistent with the combined effects of a higher Δ*V* (**Fig. 2B**) and a slower release through the small-sized outlet. Δ*V*_total_ of this configuration is more than 380% of its outlet volume, compared with only 1.4% and 10.1% for the configurations with 500 and 250 μm outlets, respectively. In principle, Δ*V* generated in 100 μm outlet configuration can be reduced by reducing the applied source pressure. However, the resulting reduction in flow rate could affect particle flushing and increase the risk of clogging during SFL process. An alternative strategy is therefore to facilitate the backward release of Δ*V* through the tubing with larger flow pathway. As a result, the required *t*_delay_ decreased substantially from 8 s in config. 3 to 2 s in config. 4 and 1 s in config. 5 when the tubing inner diameter increased from 130 μm to 250 μm and 500 μm under the condition of similar *P*_in_, flow rate and Δ*V* produced by *P*_sc_ of 2, 0.34, and 0.215 bar, respectively.

**Figure 3.**
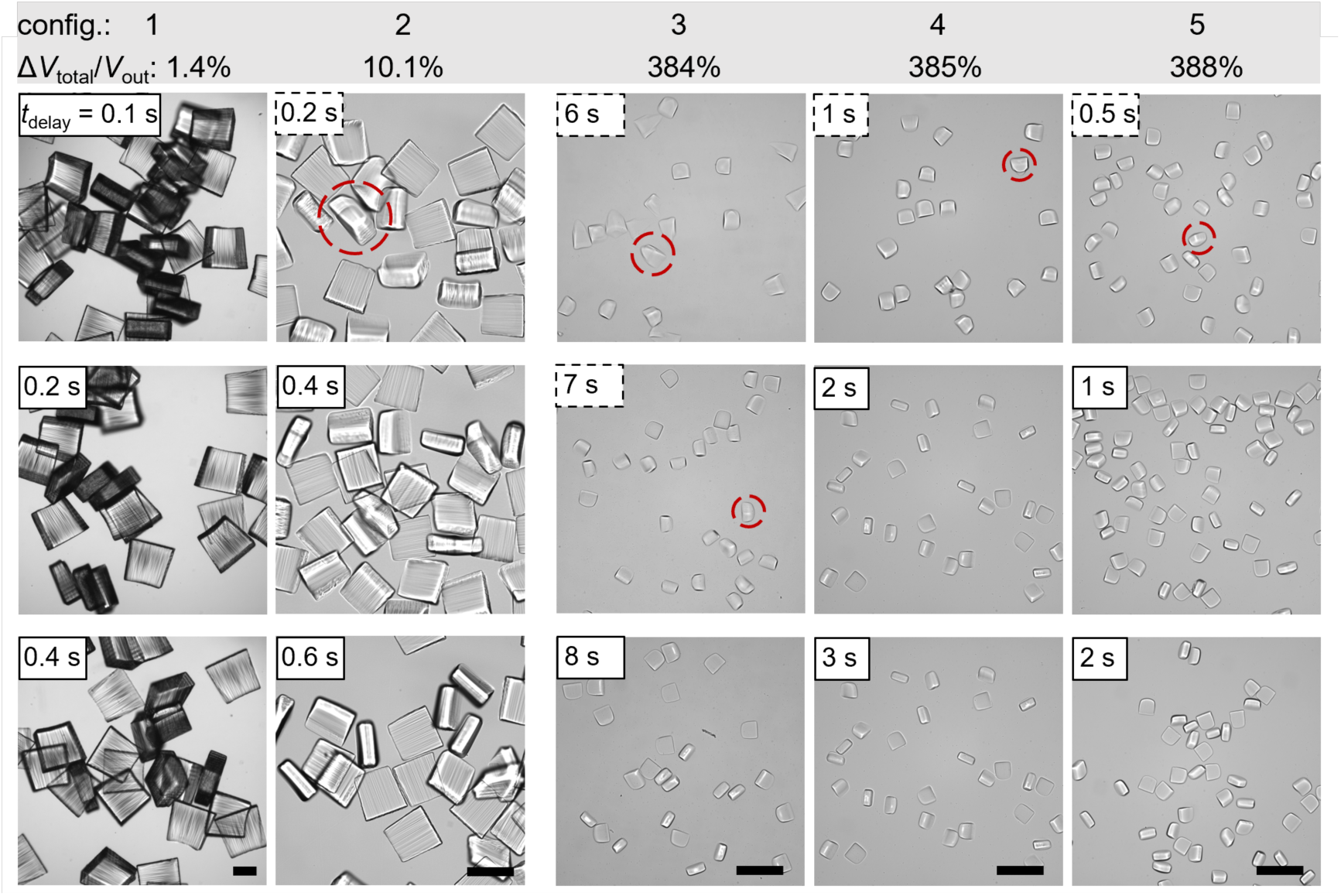
Fabrication of single-material microparticles using single-inlet configurations. Microscope images of microparticles fabricated using various single-inlet configurations at different delay times in the SFL process. Red circles highlight particles exhibiting blurred boundaries or excessive thickness, fabricated when *t*delay < *t*stopping (black dashed boxes). The corresponding fabrication parameters are provided in **Table S6**. The ratios Δ*V*total/*V*out indicated above the images provide a rough indicator for comparing flow-stopping delays across different configurations, assuming that all the Δ*V*total is released through the outlet. Scale bars: 250 μm.

To relate the simulated compliance to the observed flow-stopping delays, we estimated the characteristic discharge time of the inlet port as *τ* = *C*_in,m_·(*R*_t_ ∥ *R*_out_), where *R*_t_ ∥ *R*_out_ = *R*_t_*R*_out_/(*R*_t_ + *R*_out_) is the equivalent resistance of the two parallel release paths through the tubing and the outlet; the small inlet port resistance is neglected (**Table 2**). This lumped estimate neglects the distributed compliances of the tubing and outlet, as well as the increase of *C*_in,m_ at low pressure during discharge (Fig. 2E), and is therefore expected to underestimate *t*_stopping_. Nevertheless, it reproduces the ranking of the experimentally bracketed *t*_stopping_ across all five configurations, and *τ* lies within an order of magnitude of *t*_stopping_ in each case. In particular, it captures why config. 5, despite having the same outlet and a similar Δ*V*_total_ as configs. 3 and 4, has a *τ* more than 20-fold shorter than config. 3: the low-resistance tubing opens a fast backward release path.

**Table 2.** Estimated discharge time constant *τ* of the inlet port compliance in single-inlet configurations compared with the experimentally bracketed flow-stopping time *t*stopping.

| config. | *R*<sub>t</sub> (Pa·s/m³) | *R*<sub>out</sub> (Pa·s/m³) | *R*<sub>t</sub> ‖ *R*<sub>out</sub> (Pa·s/m³) | *C*<sub>in,m</sub> (µL/bar) | *τ* (s) | *t*<sub>stopping</sub> (s), exp. |
|---|---|---|---|---|---|---|
| 1 | 4.72×10<sup>13</sup> | 1.20×10<sup>11</sup> | 1.20×10<sup>11</sup> | 10.64 | 0.013 | ≤ 0.1 |
| 2 | 1.76×10<sup>14</sup> | 1.92×10<sup>12</sup> | 1.90×10<sup>12</sup> | 8.21 | 0.16 | 0.2–0.4 |
| 3 | 6.46×10<sup>14</sup> | 7.50×10<sup>13</sup> | 6.72×10<sup>13</sup> | 6.29 | 4.2 | 7–8 |
| 4 | 4.72×10<sup>13</sup> | 7.50×10<sup>13</sup> | 2.90×10<sup>13</sup> | 6.29 | 1.8 | 1–2 |
| 5 | 2.95×10<sup>12</sup> | 7.50×10<sup>13</sup> | 2.84×10<sup>12</sup> | 6.29 | 0.18 | 0.5–1 |

\* *R*<sub>t</sub> and *R*<sub>out</sub> from Table 1; *C*<sub>in,m</sub> is the secant compliance at the operating pressure from Fig. 2C; *τ* = *C*<sub>in,m</sub>·*R*<sub>t</sub>*R*<sub>out</sub>/(*R*<sub>t</sub> + *R*<sub>out</sub>). The experimental *t*<sub>stopping</sub> is bracketed by the longest *t*<sub>delay</sub> that produced particles with blurred boundaries and the shortest *t*<sub>delay</sub> that produced particles with well-defined boundaries (Fig. 3); for config. 1, no blurred particles were observed at the shortest tested delay of 0.1 s.

**Figure 4.**
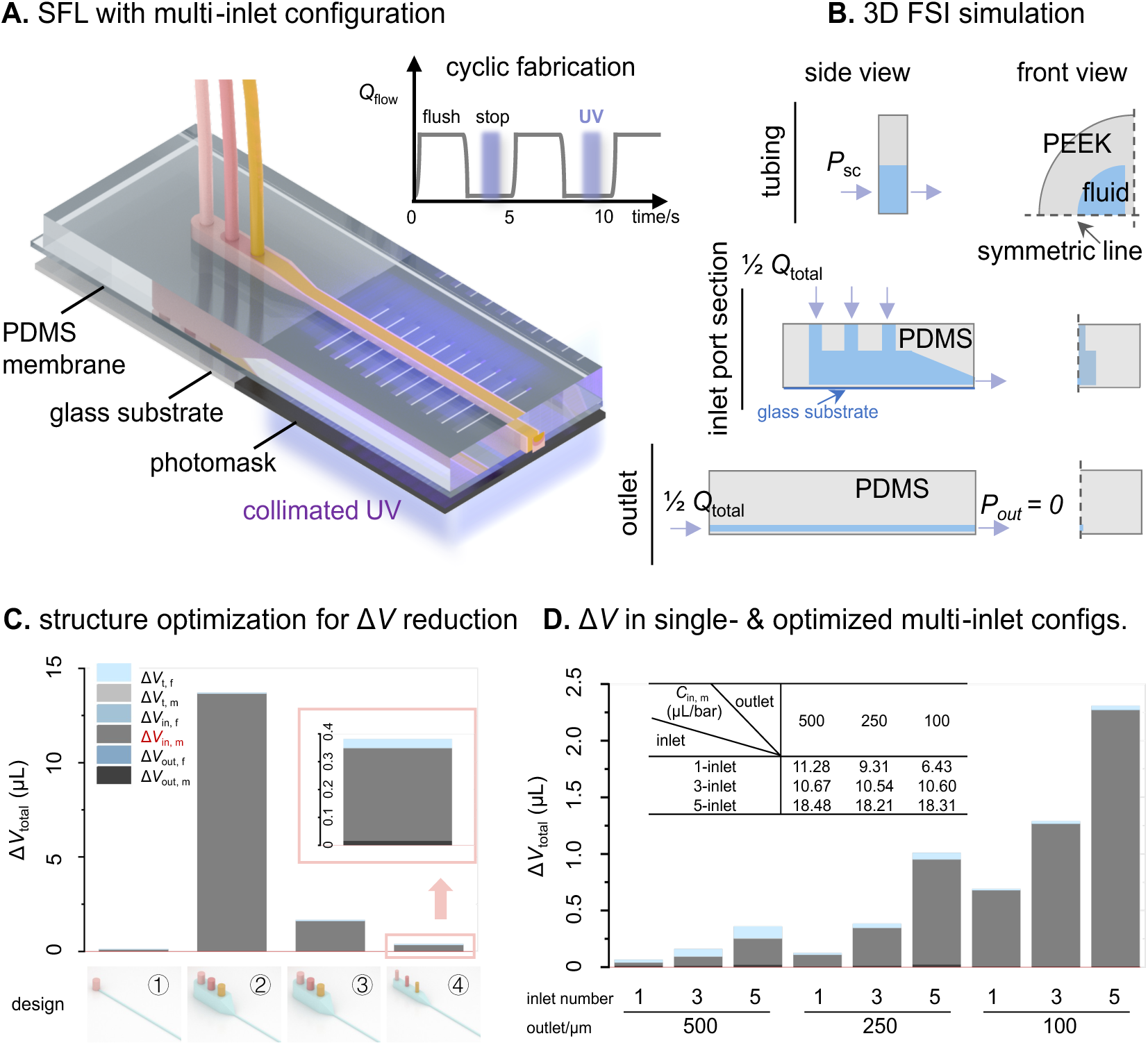
Numerical modeling and analysis of volume changes Δ*V* and hydraulic compliances *C* in multi-inlet configurations. (A) SFL using a microfluidic chip with 3-inlet configuration. Inset shows the synchronized UV exposure and flow-stop cycles of SFL process. (B) 3D models used for FSI simulations in multi-inlet configuration, including a quarter-symmetry model of the PEEK tubing, a symmetry model of the PDMS inlet port section, and a symmetry model of the PDMS outlet. Boundary conditions are provided in **Table S4**. (C) Δ*V*total and their contributions in single- and 3-inlet configurations with structural designs ①–④, at *P*sc of 1 bar applied to each tubing, with the 250 μm outlet. The inset provides an enlarged view of Δ*V*total of configuration ④, showing the effects of glass support and reduced inlet port section dimensions for Δ*V* reduction. Δ*V* data is provided in **Table S5**. (D) Δ*V*total and their contributions in single-inlet and optimized multi-inlet configurations with 1, 3, or 5 inlets and outlets of 500, 250, or 100 μm, at a source pressure of 1000 mbar (500 and 250 μm outlets) or 170 mbar (100 μm outlets, tubing of config. 4) applied to each tubing (Table S4). The inset table summarizes the corresponding *C*in, m for each configuration.

Since single-inlet configurations are limited to the fabrication of single-material microparticles, multi-inlet configurations are generally required to introduce and sculpt multiple streams for the fabrication of patterned or multi-material microparticles^33^ (**Fig. 4A**). We therefore extended our numerical modeling (**Fig. 4B**) and analysis of Δ*V* to multi-inlet configurations. As observed in single-inlet configurations, Δ*V*_in, m_ (red text, medium-gray region) remains the dominant contribution to Δ*V*_total_ in multi-inlet configurations (**Fig. 4C**). However, Δ*V*_total_ in multi-inlet configurations is substantially higher than that in the single-inlet configuration (**Fig. 4C**, designs ① and ②, 0.123 μL versus ∼13.7 μL; extrapolated from a simulation of a 3D slice of the inlet port section, Fig. S6) due to the pronouncedly higher Δ*V*_in, m_. To reduce Δ*V*_in, m_ in multi-inlet configurations, we supported the flexible PDMS membrane with a rigid glass substrate. The glass substrate effectively restricts the deformation of the PDMS membrane (**Fig. S5**) and thus significantly reduced Δ*V*_in, m_ and Δ*V*_total_ (**Fig. 4C**, designs ② and ③, ∼13.7 μL versus 1.661 μL). In addition, we also reduced the geometric dimensions of the inlet port section from 3.0 mm × 3.0 mm to 1.5 mm × 1.5 mm (together with a smaller tubing O.D. of 0.794 mm, Table S5), which further decreased Δ*V*_in, m_ (**Fig. 4C**, designs ③ and ➃, 1.661 μL versus 0.381 μL). These two optimization strategies together bring Δ*V*_total_ in multi-inlet configuration to a level comparable to that of the single-inlet configuration (**Fig. 4C**, designs ① and ➃, 0.123 μL versus 0.381 μL). **Fig. 4 D** compares Δ*V*_total_ across different single-inlet and optimized multi-inlet configurations. In general, for configurations with same outlet dimensions, Δ*V*_total_ increases with the increase of the inlet number, which is typically associated with larger dead volume of the inlet port section. Similar to the single-inlet configurations, Δ*V*_total_ in multi-inlet configurations also strongly depends on the outlet configuration. Under same inlet number conditions, configurations with 100 μm outlet consistently exhibit higher Δ*V*_total_ than those with larger outlets. *C*_in,m_ for corresponding configurations were also calculated and listed in the inset table in **Fig. 4 D**. For single-inlet configurations (without glass substrate), *C*_in,m_ decreases with the increase of the *P*_in_ (configurations with outlets of 500, 250, 100 μm) as discussed above. For multi-inlet configurations (with glass substrate), *C*_in,m_ remains approximately constant for a given number of inlets (reflecting the similar dead volume of the inlet port section), while it generally increases with the number of inlets (corresponding to the larger dead volume of the inlet port section).

We fabricated microchannels with the designs proposed in **Fig. 4C**. The experimental results of microparticle fabrication show a consistent trend with the predicted Δ*V*_total_ discussed above. As shown in **Fig. 5A**, the single-inlet configuration, which exhibits relatively low Δ*V*_total_, enabled the fabrication of particles with well-defined boundaries at a short *t*_delay_ of 0.4 s. In contrast, for the 3-inlet configuration, which exhibits substantially higher Δ*V*_total_, even a long *t*_delay_ of 12 s still resulted in continuous fibers rather than particles. Supporting the PDMS membrane with a rigid glass substrate, which significantly reduced the Δ*V*_total_, therefore enabled the formation of discrete particles at a delay of 1 s, although continuous fibers still formed at 0.4 s. With further optimized 3-inlet configuration with smaller inlet port dead volume, discrete particles with well-defined boundaries could be fabricated at a short *t*_delay_ of only 0.4 s, comparable to that of the single-inlet configuration. Note that the chips with design ④ were additionally fabricated with stiffer PDMS (5:1 base-to-curing-agent ratio, compared with 10:1 for design ③) (Section 4.1), which is not captured by the simulations (E = 580 kPa); the improvement from design ③ to ④ therefore reflects the combined effects of the reduced inlet port section, the smaller tubing, and the higher PDMS stiffness. We also fabricated optimized 3-inlet microchannels outlets in different sizes to investigate whether the designed flow profiles could be faithfully transferred into particle patterns across different particle sizes. As shown in **Fig. 5B**, for the optimized 3-inlet configurations with 500 and 250 μm outlets, the obtained flow profiles were readily transferred into the corresponding particle patterns. However, for the optimized 3-inlet configurations with a 100 μm outlet, only config. 4 successfully preserved the obtained flow profile. In config. 3, although the designed flow profile could be established during flow, it was disrupted because of the prolonged flow-stopping process. On the contrary, in config. 5, the designed flow profile could not even be established because the low hydraulic resistance of the inlet tubing was insufficient to suppress pressure fluctuations from the pressure controller, resulting in intermittent backflow.

**Figure 5.**
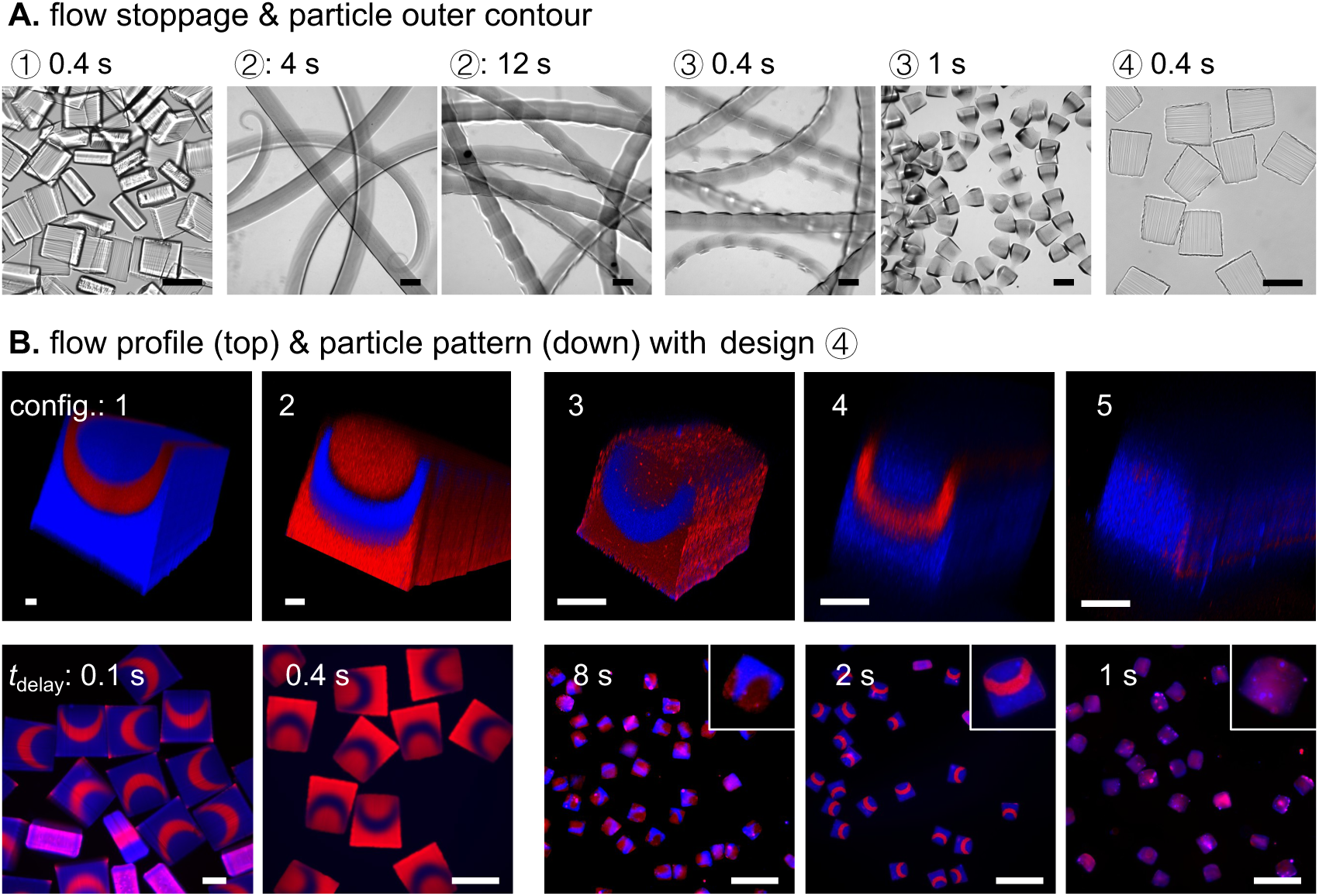
Fabrication of patterned microparticles using 3-inlet configurations. (A) Microscope images showing the effects of flow stopping delays on particle fabrication using the designs ①–④ shown in Fig. 4C (250 μm outlet) at different *t*delay; fabrication parameters as listed in Table S6 (config. 2), with only *t*delay varied. (B) Confocal microscope images of flow profiles in outlets (top) and fluorescence microscope images of the corresponding fabricated microparticles (bottom) using the optimized design ➃ shown in Fig.4C, with 500, 250, and 100 μm outlets. Fabrication parameters are listed in **Table S7**. Scale bars: 250 μm.

## 3. Conclusions

In this work, we developed a pneumatic-controlled SFL platform and systematically investigated the effects of hydrodynamic flow resistance and hydraulic compliance on flow-stopping behavior in single- and multi-inlet microfluidic configurations. Using fluid–structure interaction simulations and theoretical analysis, we identified the dominant contribution to Δ*V*_total_ and demonstrated how its generation and release can be controlled to improve microparticle fabrication.

In single-inlet configurations, deformation of the PDMS inlet port Δ*V*_in, m_ was identified as the dominant contribution to Δ*V*_total_, with its magnitude strongly influenced by the inlet pressure *P*_in_, PDMS Young’s modulus *E*, and outlet dimension, which is channel configurations in our system. By regulating the flow resistance of the tubing relative to that of the outlet channel, the generated Δ*V*_total_ could be preferentially released forward through the outlet or backward through the tubing. This strategy reduced the required flow-stopping delay from 8 to 1 s in single-inlet configuration with 100 μm outlet, enabling the fabrication of small-sized single-material microparticles with well-defined boundaries.

In multi-inlet configuration, the deformation of the PDMS inlet port section Δ*V*_in, m_ remained the dominant contribution to Δ*V*_total_, but exhibited significantly higher magnitude than that in the single-inlet configuration. Supporting the PDMS membrane beneath the inlet port section with a rigid glass substrate and reducing the dimensions of this section decreased the magnitude of Δ*V*_in, m_ and thus Δ*V*_total_ from ∼13.7 μL to 0.381 μL. The optimized 3-inlet designs enabled patterned microparticle fabrication at delay times of 0.1 and 0.4 s for the 500 and 250 μm outlets, respectively, and 2 s for the 100 μm outlet using config. 4.

Moreover, we found that the introduction of multiple inlets may change the pressure distribution, thereby modifying the relationship between the applied pressure and the total outlet velocity. In the 250 and 500 μm configurations, the vast majority of the pressure drop occurs along the tubing, effectively transforming the nominally constant pressure source into a quasi-constant flow rate source in the multi-inlet cases. In contrast, in the 100 μm configurations, the total outlet velocity becomes strongly dependent on the inlet parallelization (Fig. S7 and Supplementary Note 1).

Together, these findings establish hydrodynamic flow resistance regulation and hydraulic compliance minimization as complementary strategies for improving flow-stopping performance in SFL. By enabling the fabrication of size-tunable microparticles with well-defined boundaries and facilitating the transfer of sculpted flow profiles into particle patterns, this platform provides practical design and operating principles for reliable multi-inlet SFL systems.

## 4. Materials and Methods

### 4.1 Microchannel design and fabrication

The microchannel design and fabrication followed the procedures described in our previous work^33^. Briefly, microchannel models were designed using 3ds Max (Autodesk Inc.) and exported as SAT files for numerical modeling and STL files for 3D printing. After the microchannel molds were 3D-printed (Sonic Mini 8K printer with Aqua-Gray 8K resin, Phrozen Tech Co., Ltd.) and surface-treated (UV post-curing, oxygen plasma activation, and vapor-phase silanization with trichloro(1H,1H,2H,2H-perfluorooctyl)silane, 448931, Sigma-Aldrich), standard soft lithography with PDMS (Sylgard 184 Silicone Elastomer Kit, Dow Inc.; base-to-curing-agent mass ratio of 10:1 unless stated otherwise; cured at 65 °C for 1 h) was used to fabricate the microfluidic chips for SFL experiments. A thin PDMS membrane (100–150 μm thick, spin-coated at 500 rpm for 30 s and cured at 65 °C for 15 min) was used to seal the microchannel by oxygen plasma bonding. Specifically, a glass substrate (cover glass, 20 × 26 × 0.4 mm, Paul Marienfeld GmbH & Co. KG) was attached with a thin layer of uncured PDMS only beneath the thin PDMS membrane of the inlet port section in multi-inlet configurations, while the outlet channel was left unsupported because an additional glass layer could interfere with UV exposure and the contribution of Δ*V*_out, m_ to Δ*V*_total_ is relatively small. For the optimized 3- and 5-inlet configurations, the microfluidic chips were fabricated using a 5:1 PDMS base-to-curing-agent ratio and further heated at 90 °C for 24 h after fabrication to increase PDMS stiffness.

### 4.2 Fluid properties and solution preparation

All numerical simulations were conducted using a working fluid with a density of 987 kg·m^-3^ and a viscosity of 7.55 mPa.s, corresponding to a precursor solution consisting of 60 vol.% Poly (ethylene glycol) diacrylate (PEGDA, *M_w_*575, 437441, Sigma-Aldrich) and 40 vol.% ethanol (1.00983, Sigma-Aldrich), with an additional 5 vol.% photoinitiator (2-hydroxy-2 methylpropiophenone, Darocur 1173, 405655, Sigma-Aldrich). To visualize the particle patterns, either fluorescent dyes (50 μg/mL; red: methacryloxyethyl thiocarbamoyl rhodamine B, 23591, Polysciences, Inc.; blue: methacrylate-modified blue-emitting dye, MA-P4VB: 2-(methacryloyloxy)propyl 6-(4-methoxy-2,5-bis((E)-2-(pyridin-4-yl)vinyl)phenoxy)hexanoate, kindly provided by Dr. Sergei I. Vagin) or a fluorescent nanoparticle dispersion (5 vol.%) was added to the precursor solution.

### 4.3 Estimation of outlet velocity *v* and flow rate *Q* via numerical modeling

The Single-Phase Flow (Laminar Flow) module in COMSOL Multiphysics was used to estimate the flow velocities *v* at the outlet. The entire model, including the equivalent inlet tubing and the PDMS channel, was simulated as a union. Tubing with a diameter of 0.05 mm and lengths of 13.13, 3.57, 0.96, and 0.06 mm were used to provide flow resistances equivalent to those of tubing with a length of 600 mm and inner diameters of 0.13, 0.18, 0.25, and 0.50 mm, respectively. The boundary conditions are *P*_sc_ and *P*_out_ = 0 for the inlet of the tubing and outlet of the outlet channel, respectively. The outlet velocities obtained were plotted in **Fig. S7** to show the relationship between the outlet velocity and inlet number in various channel configurations. The outlet velocities obtained then were also converted into the corresponding flow rates *Q*_total_. Then 1/2 *Q*_total_ were used as the boundary conditions for the later Fluid–Structure Interaction (FSI) simulation of multi-inlet configuration (**Table S4**). Rough outlet velocity of single-inlet configurations listed in **Table S1** are obtained using the online microfluidic calculator.

### 4.4 Estimation of Δ*V*_m_ via numerical modeling and calculation for *C*

The FSI module in COMSOL Multiphysics was used to estimate Δ*V* of the pressurized microfluidic system. To simplify the computation, the single-inlet model was divided into two parts: i) the tubing (PEEK, Young’s modulus: 3.9 GPa; Poisson’s ratio: 0.36, 600 mm length) and ii) the PDMS channel (Young’s modulus: 280, 580, 1000 kPa, Poisson’s ratio: 0.49), comprising a 100 μm-thick bottom PDMS membrane, an inlet port with 1.5 mm diameter and 5 mm height, and an outlet with 35 mm length. Similarly, the multi-inlet model was divided into three parts: i) the tubing, ii) the inlet port section with a 100 μm-thick bottom PDMS membrane (Young’s modulus: 580 kPa, Poisson’s ratio: 0.49), and iii) the outlet channel with a 100 μm-thick bottom PDMS membrane (Young’s modulus: 580 kPa, Poisson’s ratio: 0.49). To represent the supporting glass substrate, the bottom constraint of the PDMS membrane was set as a fixed constraint. Geometric nonlinearity was included in all FSI simulations, and PDMS was modeled as a linear elastic material. For the boundary conditions used in the single-inlet and multi-inlet configurations, refer to **Tables S2 and S4**, respectively. The secant hydraulic compliances *C* corresponding to Δ*V* were calculated by dividing the magnitude of each Δ*V* by their corresponding average pressure *P*_avg_ in the respective component (**Tables S1&S3**). Mesh convergence tests are provided in Figs. S1 and S2. For design ②, for which the 3D simulation did not converge, ΔV of the inlet port section was estimated by extrapolating a simulation of a 3D slice of this section (Fig. S6).

### 4.5 Estimation of Δ*V*_f_ based on the effective bulk modulus *K*_e_

Fluid compressibility was estimated using the bulk modulus, *K*, which relates the relative volume change of a liquid to a change in subjected pressure. The bulk modulus of ethanol is 0.89 GPa at 20 °C. For PEGDA 575, the bulk modulus was estimated using *K* = *ρc*^2^, where *ρ* = 1121 kg m^−3^ and *c* = 1552 m s^−1,34^ giving *K*_PEGDA_ = 2.69 GPa at 20 °C. The bulk modulus of the 60 vol.% PEGDA–40 vol.% ethanol mixture was then estimated based on the volume fractions and bulk moduli of the two components, giving *K*_mix_ = 1.49 GPa. A study reported that the effective bulk modulus of a liquid can be ∼20-fold lower than the literature value due to dissolved air^23^. Accordingly, an effective bulk modulus *K*_e_ of ∼ 0.075 GPa was assumed for the precursor solution. Then Δ*V* were calculated using │Δ*V*│ = │-*V*_0_*P*_avg_/*K*_e_*│*, where *V*_0_ is the fluid dead volume, *P*_avg_ is the average pressure.

### 4.6 Stop-flow lithography

A standard SFL procedure was used to fabricate microparticles, as detailed in our previous work^13–16,33^. Briefly, a customized python script was used to automate the system by sending the synchronized signals to the pressure controllers (LineUp Flow EZ 7000, Fluigent Deutschland GmbH) and the UV light unit (S2000 Elite, Excelitas Canada Inc., coupled with a collimator and operated at 100% intensity), thereby achieving periodic flow-stop cycles of the precursor flow with UV exposure precisely timed to the flow-stopping phase through photomask. The photomasks were customized from Zitzmann Plot GmbH. PEEK tubing (Analytics-Shop; O.D. 1/16 inch for single-inlet and design ①–③ chips, 1/32 inch for design ➃ chips) connected the reservoirs to the chip. The fabricated particles/fibers were collected from the opened outlet channel, thoroughly washed three times to remove uncured materials, and then stored in either ethanol or a 0.1% Pluronic^®^ F-127 (P2443, Sigma-Aldrich) aqueous solution for subsequent imaging.

## Supplementary Material

The Supplementary Information (PDF) contains Tables S1–S7, Figures S1–S7, and Supplementary Note 1. Supporting Data 1 (Excel) and Numerical Models 1–3 (COMSOL Multiphysics .mph files) are provided as separate files.

## Acknowledgment

Y.Z. gratefully acknowledges the China Scholarship Council for the doctoral fellowship. We thank Dr. Sergei I. Vagin for kindly providing the blue-emitting dye MA-P4VB.

## Data Availability Statement

The data that support the findings of this study are available in the Supplementary Information and the accompanying supplementary files: Supporting Data 1 (Excel workbook with the simulated volume changes and hydraulic compliances underlying Figs. 2 and 4, Table 2, and Tables S3 and S5), and Numerical Models 1–3 (COMSOL Multiphysics 6.3 model files of the PEEK tubing, the single-inlet PDMS channel, and the multi-inlet PDMS channel, respectively). Additional data are available from the corresponding author upon reasonable request.

## Declaration of AI-Assisted Technologies

During the preparation of this manuscript, the authors used generative AI tools based on large language models to proofread the text for grammatical and typographical errors, to check the consistency of numerical values, cross-references, and figure captions, and to suggest wording for selected passages. All suggestions were reviewed, verified, and edited by the authors, who take full responsibility for the content of the publication.

## Supplementary Information

**Table S1.**
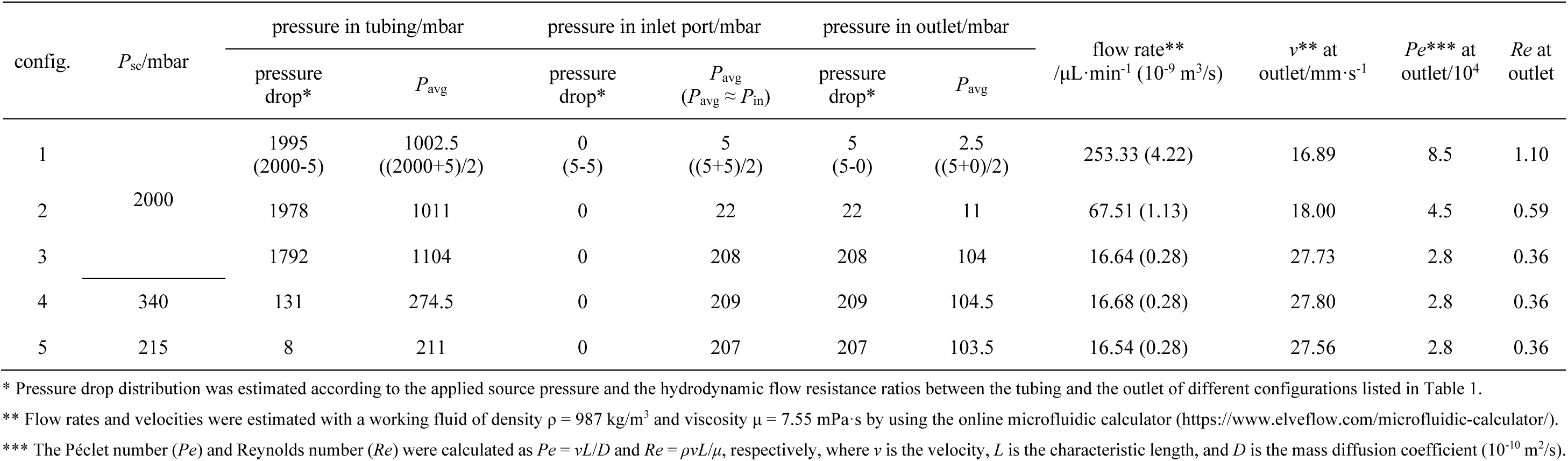
Flow characteristics estimation for the single-inlet configurations.

| config. | *P*<sub>sc</sub>/mbar | pressure in tubing/mbar |  | pressure in inlet port/mbar |  | pressure in outlet/mbar |  | flow rate**<br>/µL·min<sup>-1</sup> (10<sup>-9</sup> m<sup>3</sup>/s) | *v*\*\* at<br>outlet/mm·s<sup>-1</sup> | *Pe*\*\*\* at<br>outlet/10<sup>4</sup> | *Re* at<br>outlet |
|---|---|---|---|---|---|---|---|---|---|---|---|
|  |  | pressure<br>drop* | *P*<sub>avg</sub> | pressure<br>drop* | *P*<sub>avg</sub><br>(*P*<sub>avg</sub> ≈ *P*<sub>in</sub>) | pressure<br>drop* | *P*<sub>avg</sub> |  |  |  |  |
| 1 | 2000 | 1995<br>(2000-5) | 1002.5<br>((2000+5)/2) | 0<br>(5-5) | 5<br>((5+5)/2) | 5<br>(5-0) | 2.5<br>((5+0)/2) | 253.33 (4.22) | 16.89 | 8.5 | 1.10 |
| 2 |  | 1978 | 1011 | 0 | 22 | 22 | 11 | 67.51 (1.13) | 18.00 | 4.5 | 0.59 |
| 3 |  | 1792 | 1104 | 0 | 208 | 208 | 104 | 16.64 (0.28) | 27.73 | 2.8 | 0.36 |
| 4 | 340 | 131 | 274.5 | 0 | 209 | 209 | 104.5 | 16.68 (0.28) | 27.80 | 2.8 | 0.36 |
| 5 | 215 | 8 | 211 | 0 | 207 | 207 | 103.5 | 16.54 (0.28) | 27.56 | 2.8 | 0.36 |

\* Pressure drop distribution was estimated according to the applied source pressure and the hydrodynamic flow resistance ratios between the tubing and the outlet of different configurations listed in Table 1.
\*\* Flow rates and velocities were estimated with a working fluid of density ρ = 987 kg/m<sup>3</sup> and viscosity µ = 7.55 mPa·s by using the online microfluidic calculator (https://www.elveflow.com/microfluidic-calculator/).
\*\*\* The Péclet number (*Pe*) and Reynolds number (*Re*) were calculated as *Pe* = *vL*/*D* and *Re* = *ρvL*/*µ*, respectively, where *v* is the velocity, *L* is the characteristic length, and *D* is the mass diffusion coefficient (10<sup>-10</sup> m<sup>2</sup>/s).

**Table S2.** Simulation boundary conditions for single-inlet configurations (corresponding to Fig. 2B).

| config. | tubing (600 mm) |  |  |  |  | PDMS channel<br>comprising the inlet port and outlet channel |  |
|---|---|---|---|---|---|---|---|
|  | pressure drop/(mbar/0.1 mm) | first 0.1-mm segment |  | last 0.1-mm segment |  |  |  |
|  |  | *P*<sub>sc</sub> /mbar | outlet/mbar | inlet/mbar | outlet/mbar | *P*<sub>in</sub>/mbar | *P*<sub>out</sub>/mbar |
| 1 | 0.3325<br>((2000-5)/6000) | 2000 | 2000-0.3325 | 5+0.3325 | 5 | 5 | 0 |
| 2 | 0.3297 | 2000 | 2000-0.3297 | 22+0.3297 | 22 | 22 | 0 |
| 3 | 0.2987 | 2000 | 2000-0.2987 | 208+0.2987 | 208 | 208 | 0 |
| 4 | 0.0218 | 340 | 340-0.0218 | 209+0.0218 | 209 | 209 | 0 |
| 5 | 0.0013 | 215 | 215-0.0013 | 207+0.0013 | 207 | 207 | 0 |

**Table S3.** Volume changes Δ*V* estimation in single-inlet configurations (corresponding to Fig. 2B).

| config. | *P*<sub>sc</sub> /mbar | Δ*V*/µL |  |  |  |  |  | Δ*V*<sub>total</sub>/µL | *V*<sub>out</sub>/µL | Δ*V*<sub>total</sub>/*V*<sub>out</sub> |
|---|---|---|---|---|---|---|---|---|---|---|
|  |  | Δ*V*<sub>t, m</sub> | Δ*V*<sub>in, m</sub> | Δ*V*<sub>out, m</sub> | Δ*V*<sub>t, f</sub> | Δ*V*<sub>in, f</sub> | Δ*V*<sub>out, f</sub> |  |  |  |
| 1 | 2000 | 0.0012 | 0.0532 | 0.0306 | 0.0394 | 0.0001 | <0.0001 | 0.125 | 8.75 | 1.42% |
| 2 |  | 0.0010 | 0.1807 | 0.0186 | 0.0206 | 0.0003 |  | 0.221 | 2.1875 | 10.11% |
| 3 |  | 0.0007 | 1.3083 | 0.0197 | 0.0117 | 0.0025 |  | 1.343 | 0.35 | 383.69% |
| 4 | 340 | <0.0001 | 1.3143 | 0.0198 | 0.0108 | 0.0025 |  | 1.347 |  | 384.97% |
| 5 | 215 | 0.0024 | 1.3013 | 0.0196 | 0.0331 | 0.0024 |  | 1.359 |  | 388.23% |

**Table S4.** Simulation boundary conditions for multi-inlet configurations in COMSOL simulations (corresponding to Fig. 4D).

| config. | tubing (600 mm) |  |  |  |  | inlet port section |  | outlet channel |  |
|---|---|---|---|---|---|---|---|---|---|
|  | pressure drop/(mbar/0.1 mm) | first 0.1-mm segment |  | last 0.1-mm segment |  |  |  |  |  |
|  |  | *P*<sub>sc</sub> /mbar | outlet/mbar | inlet/mbar | outlet/mbar | 1/2\*1/n\**Q*<sub>total</sub>/m<sup>3</sup>·s<sup>-1</sup><br>(per inlet port) | outlet/mbar | 1/2\**Q*<sub>total</sub>/m<sup>3</sup>·s<sup>-1</sup> | *P*<sub>out</sub>/mbar |
| 1-inlet 500 µm outlet | 0.1663<br>((1000-2.5)/6000) | 1000 | 1000-0.1663 | 2.5+0.1663 | 2.5 | *P*<sub>in</sub>: 2.5 mbar | / | / | 0 |
| 3-inlet 500 µm outlet | 0.1654 | 1000 | 1000-0.1654 | 7.63+0.1654 | 7.63 | 1/2*1/3*6.25e-9 | 7.50 | 1/2*6.25e-9 | 0 |
| 5-inlet 500 µm outlet | 0.1646 | 1000 | 1000-0.1646 | 12.54+0.1646 | 12.54 | 1/2*1/5*10.30e-9 | 12.34 | 1/2*10.30e-9 | 0 |
| 1-inlet 250 µm outlet | 0.1648 | 1000 | 1000-0.1648 | 11+0.1646 | 11 | *P*<sub>in</sub>: 11 mbar | / | / | 0 |
| 3-inlet 250 µm outlet | 0.1614 | 1000 | 1000-0.1614 | 31.61+0.1614 | 31.61 | 1/2*1/3*1.66e-9 | 31.49 | 1/2*1.66e-9 | 0 |
| 5-inlet 250 µm outlet | 0.1582 | 1000 | 1000-0.1582 | 50.97+0.1582 | 50.97 | 1/2*1/5*2.71e-9 | 50.84 | 1/2*2.71e-9 | 0 |
| 1-inlet 100 µm outlet | 0.0109 | 170 | 170-0.0109 | 104.5+0.0109 | 104.5 | *P*<sub>in</sub>: 104.5 mbar | / | / | 0 |
| 3-inlet 100 µm outlet | 0.0085 | 170 | 170-0.0085 | 118.93+0.0085 | 118.93 | 1/2*1/3*0.19e-9 | 118.69 | 1/2*0.19e-9 | 0 |
| 5-inlet 100 µm outlet | 0.0077 | 170 | 170-0.0077 | 123.52+0.0077 | 123.52 | 1/2*1/5*0.20e-9 | 123.52 | 1/2*0.20e-9 | 0 |

\* The inlet flow rate boundary conditions for the outlet segments were determined by the velocities obtained from the corresponding full multi-inlet configuration model simulated using only laminar flow module (**Fig.S7**). The pressure measured at the inlets of outlet channel segments were then used as the outlet pressure boundary conditions for inlet port segments, and similarly the pressure measured at the inlets of inlet port segments were then used to estimate the pressure boundary conditions for tubing segments.

**Table S5.** Volume changes Δ*V* estimation in multi-inlet configurations (corresponding to Fig. 4C).

| channel design | *P*<sub>sc</sub> /mbar | pressure drop/mbar |  | Δ*V*/µL |  |  |  |  |  | Δ*V*<sub>total</sub>/µL | Δ*V*<sub>total</sub>/*V*<sub>out</sub> |
|---|---|---|---|---|---|---|---|---|---|---|---|
|  |  | tubing | PDMS channel | Δ*V*<sub>t, m</sub> \* | Δ*V*<sub>in, m</sub> | Δ*V*<sub>out, m</sub> | Δ*V*<sub>t, f</sub> | Δ*V*<sub>in, f</sub> | Δ*V*<sub>out, f</sub> |  |  |
| ① | 1000 | 989 | 11 | 0.0005 | 0.1024 | 0.0093 | 0.0103 | 0.0001 | <0.0001 | 0.123 | 5.60% |
| ② ** |  |  | 32 | 0.0014 | 13.6406 | 0.0260 | 0.0309 | 0.0043 |  | 13.703 | 626.43% |
| ③ |  |  |  |  | 1.608 | 0.0160 |  |  |  | 1.661 | 75.91% |
| ④ |  |  |  | 0.0014 | 0.3320 |  |  | 0.0009 |  | 0.381 | 17.43% |

\* The tubing O.D. was 1.588 mm for cases ①, ②, and ③, 0.794 mm for case ④.
\*\* Due to the strong fluid–structure interaction resulting from the absence of a glass substrate, the 3D numerical simulation failed to converge. Consequently, channel compliance was evaluated using a 3D slice model of the channel cross-section, which yielded deformation rates of 13.38% (Figure S6) for the inlet port section.

**Table S6.** Fabrication parameters for blank microparticle using single-inlet configurations.

| Config. | expected particle size side length × side length × thickness /mm | applied pressure/mbar | Photomask |  | exposure time/s | delay time/s | flush time/s | throughput/particles·h<sup>-1</sup> |
|---|---|---|---|---|---|---|---|---|
|  |  |  | slit-gap size/µm | slit number |  |  |  |  |
| 1 | 0.5*0.5*0.2 | 2000 | 200-200 | 66 | 0.6 | 0.1 | 5 | 4.17×10<sup>4</sup> |
| 2 | 0.25*0.25*0.1 |  | 100-200 | 88 | 0.6 | 0.4 | 3 | 7.92×10<sup>4</sup> |
| 3 | 0.1*0.1*0.05 |  | 50-200 | 105 | 0.4 | 8 | 3 | 3.32×10<sup>4</sup> |
| 4 |  | 340 |  |  |  | 2 |  | 7.00×10<sup>4</sup> |
| 5 |  | 215 |  |  |  | 1 |  | 8.59×10<sup>4</sup> |

**Table S7.** Fabrication parameters for patterned microparticle using 3-inlet configurations.

| Config. | expected particle size side length × side length × thickness /mm | applied pressure at each tubing/ mbar* | Photomask |  | exposure time/s | delay time/s | flush time/s | throughput/particles·h<sup>-1</sup> |
|---|---|---|---|---|---|---|---|---|
|  |  |  | slit-gap size/µm | slit number |  |  |  |  |
| 1 | 0.5*0.5*0.2 | 1000 | 200-200 | 66 | 0.6 | 0.1 | 5 | 4.17×10<sup>4</sup> |
| 2 | 0.25*0.25*0.1 |  | 100-200 | 88 | 0.6 | 0.4 | 4 | 6.34×10<sup>4</sup> |
| 3 | 0.1*0.1*0.05 |  | 50-200 | 105 | 0.4 | 8 | 5 | 2.82×10<sup>4</sup> |
| 4 |  | 340 |  |  |  | 2 |  | 5.11×10<sup>4</sup> |
| 5 |  | 215 |  |  |  | 1 |  | 5.91×10<sup>4</sup> |

**Figure S1.**
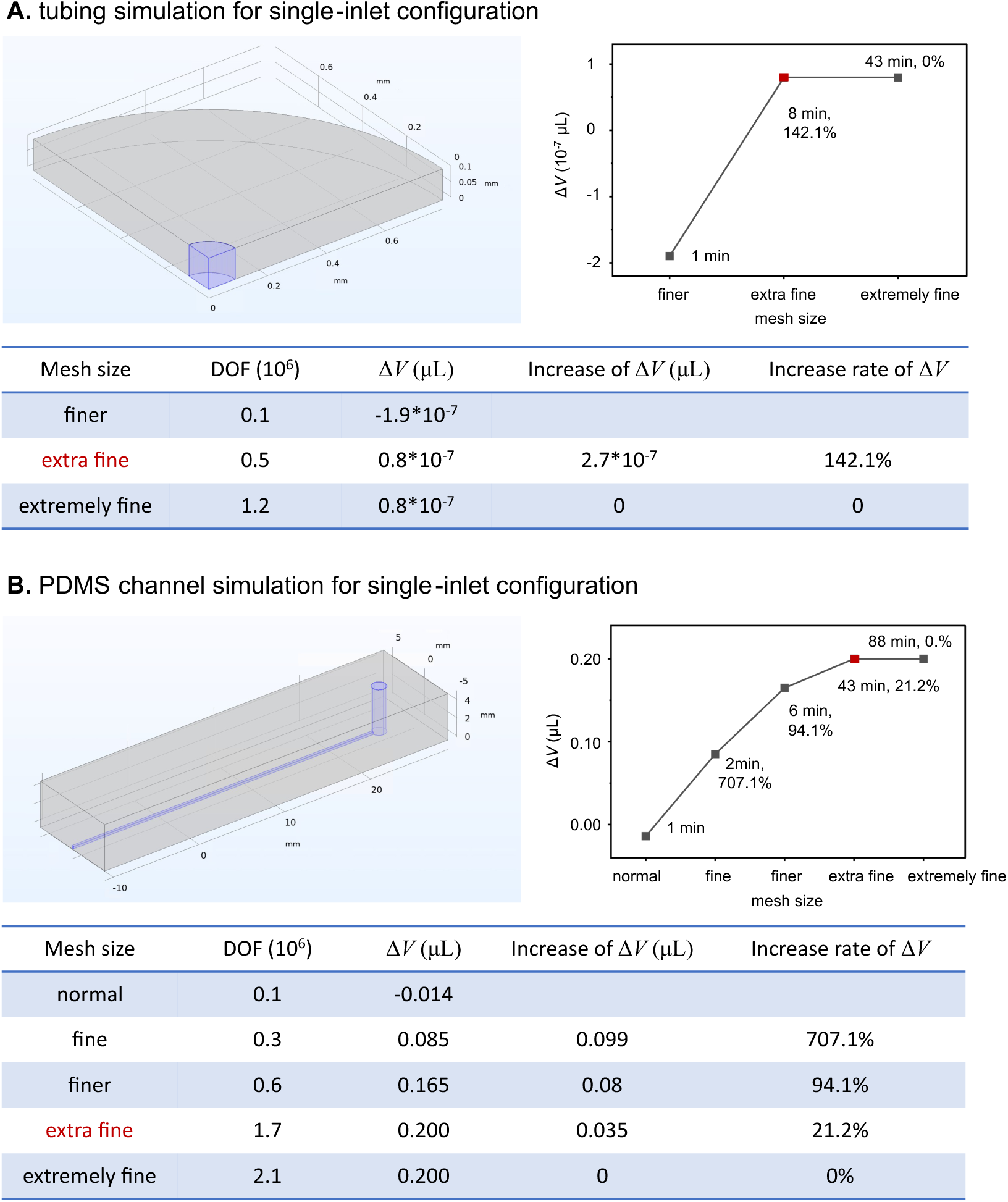
Mesh convergence test for (A) tubing and (B) PDMS channel of single-inlet configuration. The effects of mesh refinement on the volume change (Δ*V*), degrees of freedom (DOF), and computational time were evaluated. The mesh sizes highlighted in red indicate the selected mesh resolutions for subsequent simulations, providing a balance between numerical accuracy and computational efficiency.

**Figure S2.**
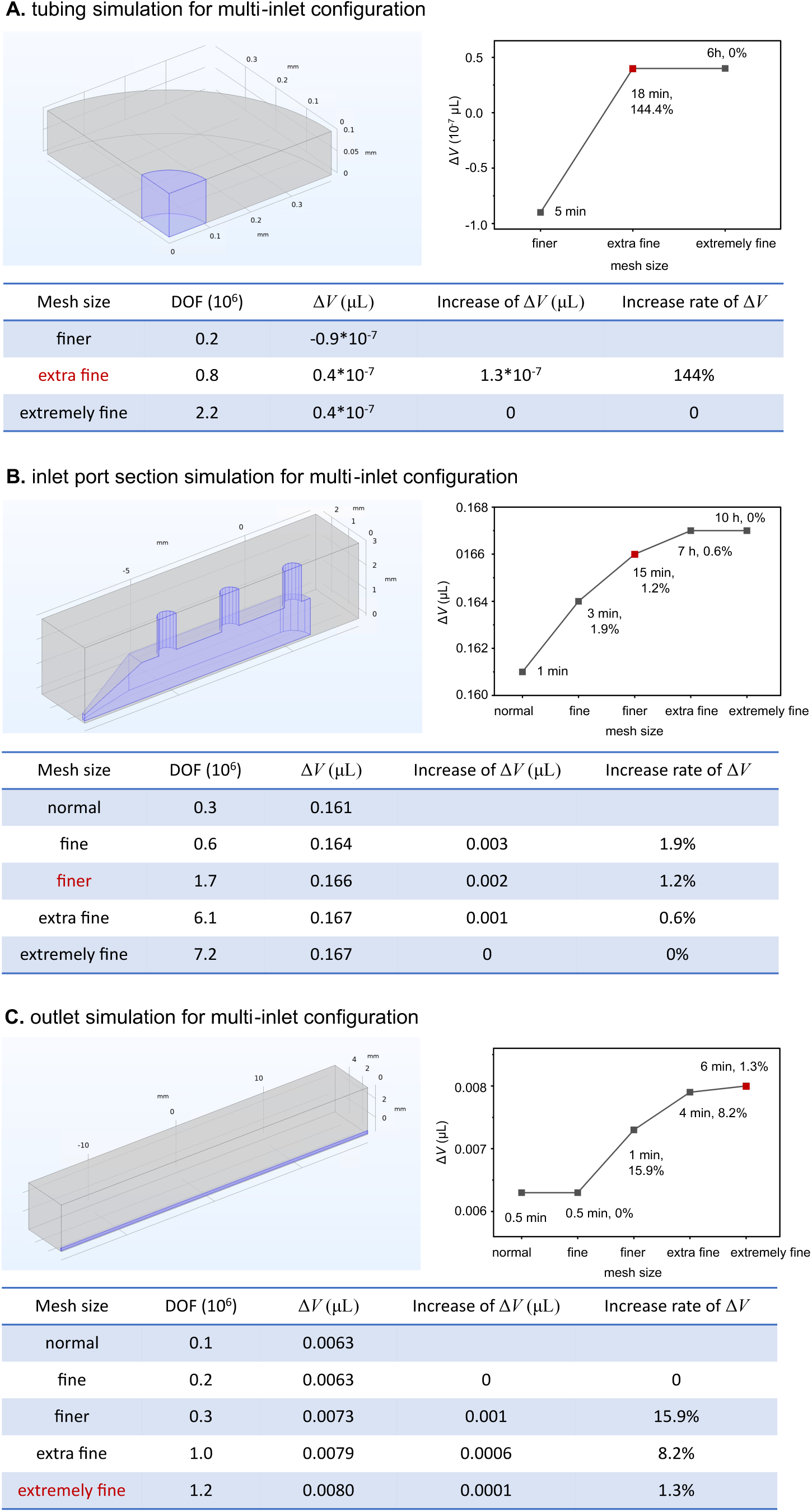
Mesh convergence test for (A) tubing, (B) inlet port section, and (C) outlet of multi-inlet configuration.

**Figure S3.**
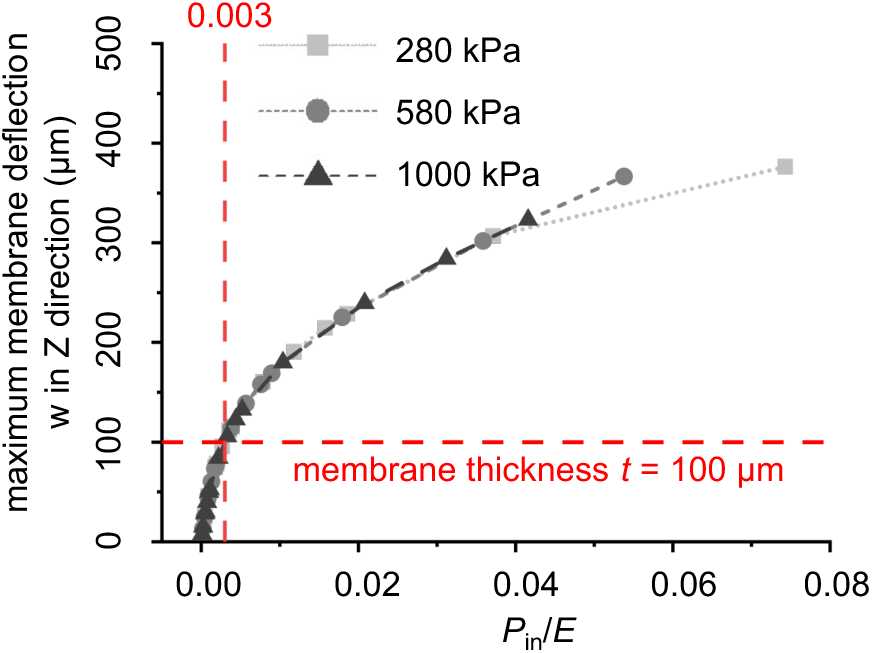
*w*–*P*_in_/*E* curves showing the normalized increase of maximum membrane deflection *w* in the z-direction with *P*_in_/*E* for three Young’s moduli. When *P*_in_/*E* exceeds 0.003, the maximum deflection w exceeds the membrane thickness *t* (100 μm); this corresponds to *P*_in_ of approximately 0.0084, 0.0174, and 0.030 bar for Young’s moduli of 280, 580, and 1000 kPa, respectively.

**Figure S4.**
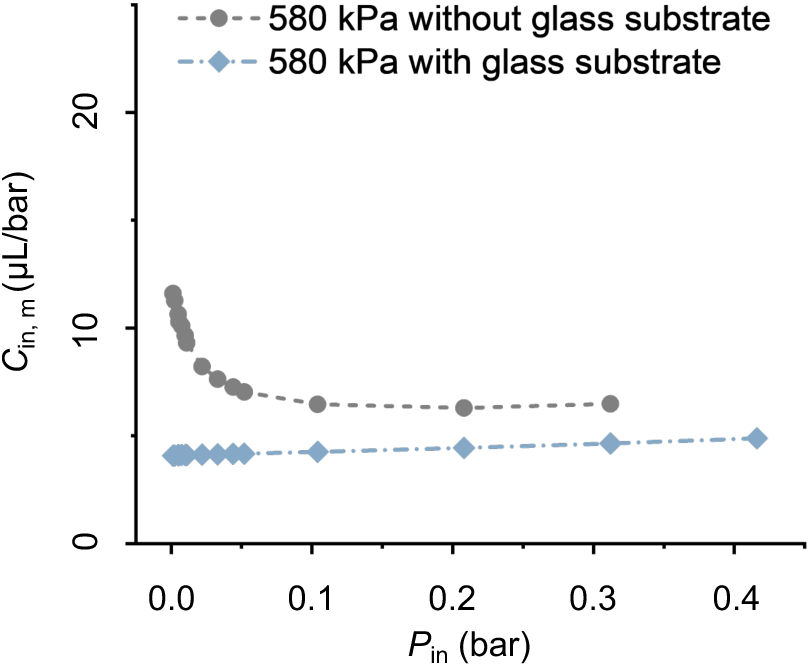
*C*_in, m_–*P*_in_ curves obtained using single-inlet configurations with and without glass substrate. Two curves indicate the distinct deforming behaviors of the inlet port under pressurized conditions.

**Figure S5.**
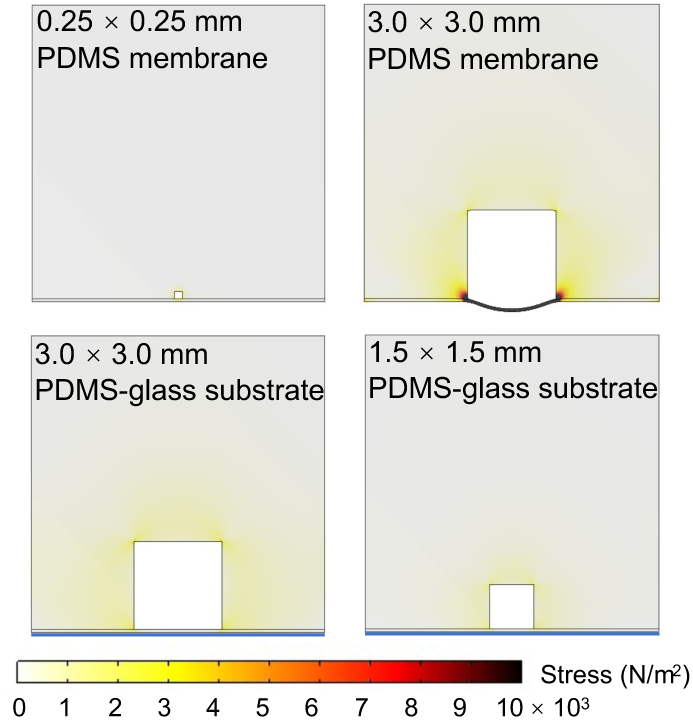
Cross-sectional deformation simulation of microchannels in different dimensions and with or without a supporting rigid glass substrate.

**Figure S6.**
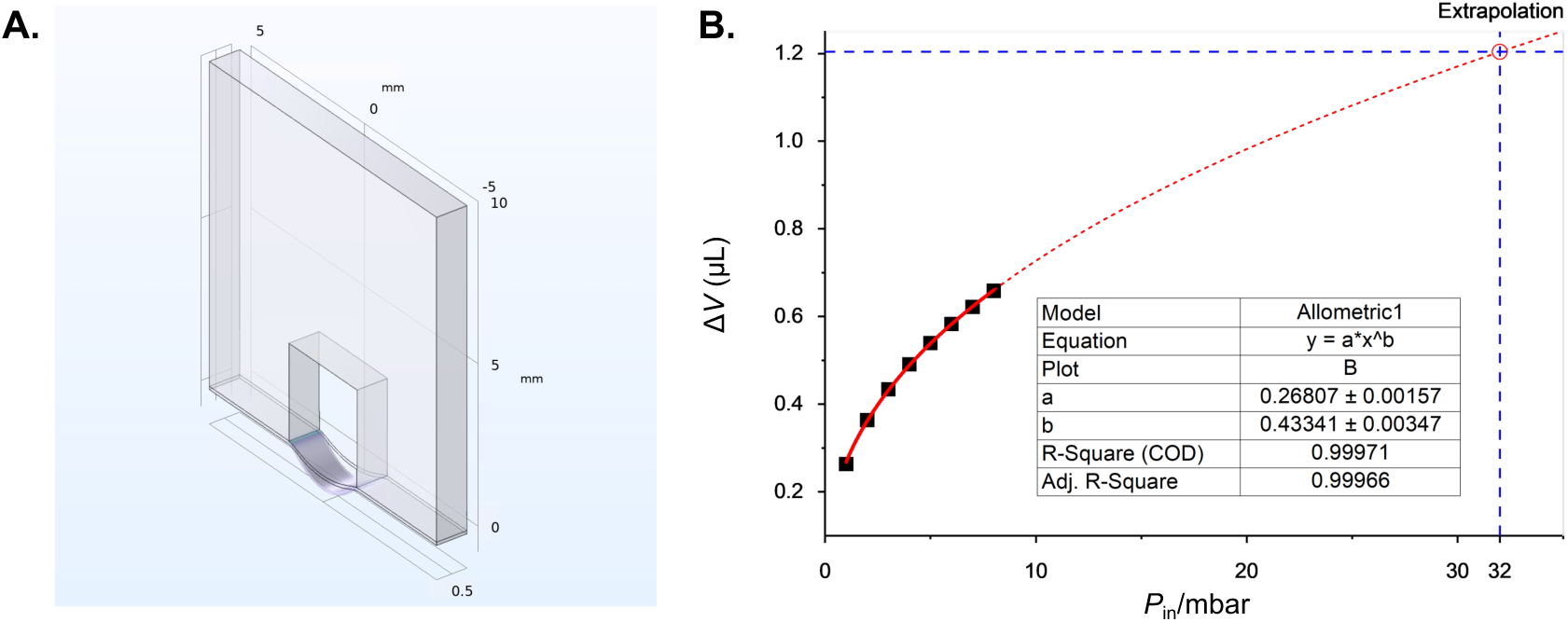
Volume change estimation using a simplified 3D slice model. Due to excessive deformation, the original 3D model failed to converge at 32 mbar, preventing direct estimation of the volume change. A simplified 3D slice model (a) was therefore employed to evaluate the pressure-dependent volume response. The simulated data (black squares) were fitted using a power-law function (solid red line, R² = 0.99971), and the resulting relationship was extrapolated to 32 mbar (dashed red line). The open circle indicates the predicted volume change of 1.204 μL at 32 mbar, corresponding to a relative volume increase of approximately 13.38% compared with the initial volume of 9 μL. Assuming the same relative volume change in the original 3D model, the volume change at 32 mbar was estimated to be 13.6406 μL, based on an initial 3D volume of 101.948 μL.

**Figure S7.**
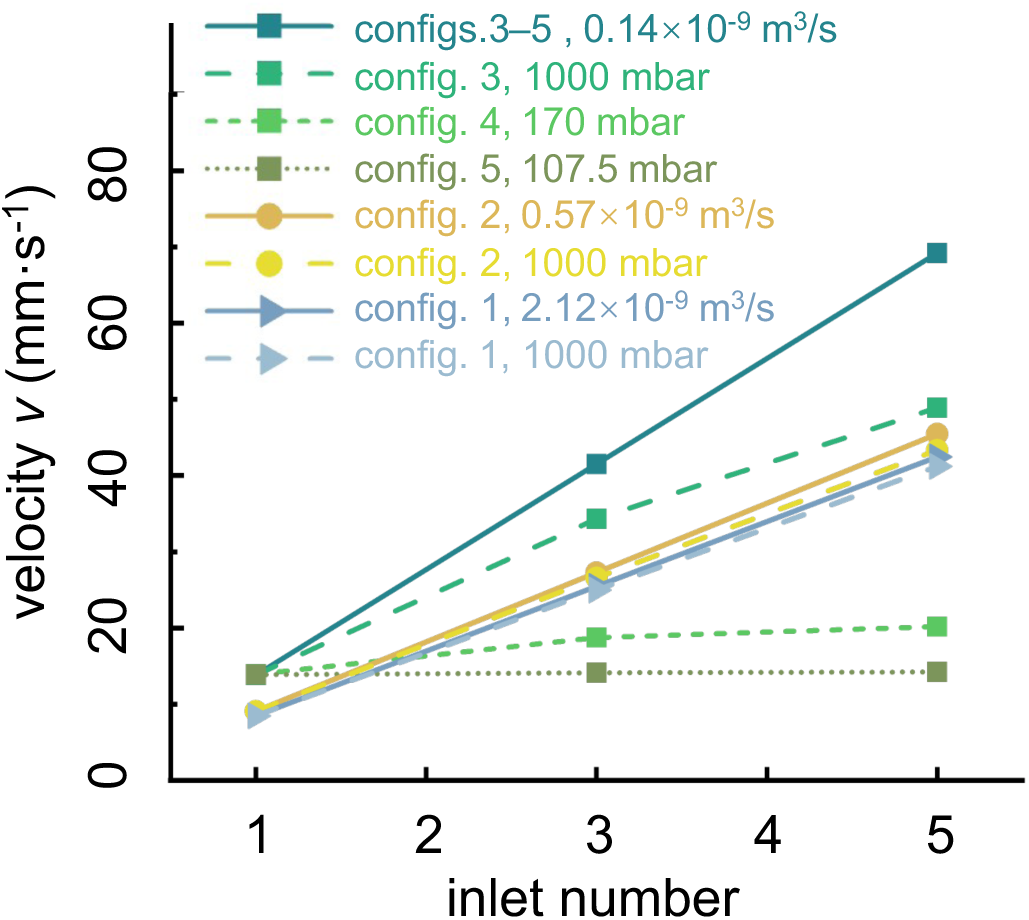
Outlet velocity *v* estimation under constant pressure *P* and flow rate *Q* boundary conditions for single- and optimized multi-inlet configurations (corresponding to Table S4 and Fig. 4D). Details are discussed in Supplementary Note 1.

**Supplementary Note 1:** Flow rate, or the corresponding outlet flow velocity, is a critical parameter in SFL, as it is closely related to the pressure required to efficiently flush the cured particles out of the microchannel between the fabrication cycles. In single-inlet configurations, applying a constant source pressure *P*_sc_ applied at the tubing produces a corresponding constant flow rate *Q* throughout the microfluidic circuit and, consequently, a constant outlet velocity *v*. The same *v* can also be obtained by directly prescribing the corresponding *Q*. For example, in single-inlet configuration with 500 μm outlet, both a pressure boundary condition of 1000 mbar and a flow rate boundary condition of 2.12 × 10^-9^ m^3^/s can produce the identical outlet velocity of 8.49 mm·s^-1^. In multi-inlet configurations with 500 μm outlet, because the flow resistance of the tubing is much higher than that of the PDMS channel, the vast majority of the *P*_sc_ is distributed at the tubing, and thus the corresponding flow rate generated by each tubing under pressure boundary condition is relatively insensitive to the PDMS channel flow resistance. This effectively transforms the nominally constant pressure source into a quasi-constant flow rate source. For example, in 500 μm outlet configurations with 3 and 5 inlets, applying a pressure of 1000 mbar at each tubing produces outlet velocities of 24.99 mm·s^-1^ (≈ 3 × 8.49 mm·s^-1^) and 41.20 mm·s^-1^ (≈ 5 × 8.49 mm·s^-1^), respectively, which are approximately the velocities obtained under flow rate boundary condition of 2.12 × 10^-9^ m^3^/s applied at each tubing in 3-and 5-inlet configurations (blueish curves of config.1). Similar situation happened in the multi-inlet configurations with 250 μm outlet (yellowish curves of config.2).

The approximation discussed above remains valid until parallelization of the tubing, analogous to resistors connected in parallel in an electrical circuit, sufficiently decrease their effective flow resistance and eliminates the dominance over the PDMS channel flow resistance. For example, in multi-inlet configurations with 100 μm outlet, the outlet velocities produced under the constant *P* and *Q* boundary conditions used in single-inlet case are not approximate, and the difference between them increases with the number of the parallel tubing (greenish curves of configs.3–5). In the extreme case of config. 5, where the flow resistance of the tubing is much lower than that of the PDMS channel, the multi-inlet configuration obtains a similar outlet velocity *V* to that of the single-inlet configuration when the same source pressure 107.50 mbar (the 1000 mbar-equivalent source pressure of config. 5, Fig. S7) is applied at each tubing.

In summary, for the 250 and 500 μm outlet configurations, distributing the total inlet pressure *P* that equals the source pressure used in the corresponding single-inlet system among multiple parallel tubing can result in an approximately identical flow velocity, whereas for 100 μm outlet like config. 5, applying the same source pressure *P* used in the single-inlet case to each parallel tubing leads to an approximately identical outlet velocity.

## References

1. Park, J., Destgeer, G., Afzal, M. & Sung, H. J. Acoustofluidic generation of droplets with tunable chemical concentrations. Lab Chip 20, 3922–3929 (2020).

2. Zou, Y. et al. Interfacial Complexation Induced Controllable Fabrication of Stable Polyelectrolyte Microcapsules Using All-Aqueous Droplet Microfluidics for Enzyme Release. ACS Appl. Mater. Interfaces 11, 21227–21238 (2019).

3. Yang, Y., Vagin, S. I., Rieger, B. & Destgeer, G. Fabrication of Crescent Shaped Microparticles for Particle Templated Droplet Formation. Macromol. Rapid Commun. 45, 2300721 (2024).

4. Akartuna, I., Aubrecht, D. M., Kodger, T. E. & Weitz, D. A. Chemically induced coalescence in droplet-based microfluidics. Lab Chip 15, 1140–1144 (2015).

5. Park, J. et al. Acoustothermal tweezer for droplet sorting in a disposable microfluidic chip. Lab Chip 17, 1031–1040 (2017).

6. Choi, S. & Park, J.-K. Continuous hydrophoretic separation and sizing of microparticles using slanted obstacles in a microchannel. Lab Chip 7, 890 (2007).

7. Destgeer, G. et al. Microchannel Anechoic Corner for Size-Selective Separation and Medium Exchange via Traveling Surface Acoustic Waves. Anal. Chem. 87, 4627–4632 (2015).

8. Tasadduq, B., Lam, W., Alexeev, A., Sarioglu, A. F. & Sulchek, T. Enhancing size based size separation through vertical focus microfluidics using secondary flow in a ridged microchannel. Sci Rep 7, 17375 (2017).

9. Magnusson, C., Rezayati Charan, M. & Augustsson, P. Two-Step Acoustic Cell Separation Based on Cell Size and Acoustic Impedance─toward Isolation of Viable Circulating Tumor Cells. Anal. Chem. 97, 2120–2126 (2025).

10. Mudugamuwa, A. et al. Periodic Flows in Microfluidics. Small 20, 2404685 (2024).

11. Mohammed, M. et al. Studying the Response of Aortic Endothelial Cells under Pulsatile Flow Using a Compact Microfluidic System. Anal. Chem. 91, 12077–12084 (2019).

12. Dendukuri, D., Gu, S. S., Pregibon, D. C., Hatton, T. A. & Doyle, P. S. Stop-flow lithography in a microfluidic device. Lab Chip 7, 818 (2007).

13. Destgeer, G., Ouyang, M., Wu, C.-Y. & Di Carlo, D. Fabrication of 3D concentric amphiphilic microparticles to form uniform nanoliter reaction volumes for amplified affinity assays. Lab Chip 20, 3503–3514 (2020).

14. Destgeer, G., Ouyang, M. & Di Carlo, D. Engineering Design of Concentric Amphiphilic Microparticles for Spontaneous Formation of Picoliter to Nanoliter Droplet Volumes. Anal. Chem. 93, 2317–2326 (2021).

15. Akhtar, M. U., Sahin, M. A., Werner, H. & Destgeer, G. Fabrication of Size-Coded Amphiphilic Particles with a Configurable 3D-Printed Microfluidic Device for the Formation of Particle-Templated Droplets. Adv Materials Technologies 9, 2301967 (2024).

16. Song, X. et al. Tunable Picoliter-Scale Dropicle Formation Using Amphiphilic Microparticles with Patterned Hydrophilic Patches. Advanced Science 12, 2411014 (2025).

17. Leng, L., McAllister, A., Zhang, B., Radisic, M. & Günther, A. Mosaic Hydrogels: One-Step Formation of Multiscale Soft Materials. Advanced Materials 24, 3650–3658 (2012).

18. Carbonell, C., Linares-Moreau, M., Borisov, S. M. & Falcaro, P. Multimaterial Digital-Light Processing of Metal-Organic Framework (MOF) Composites: A Versatile Tool for the Rapid Microfabrication of MOF-Based Devices. Advanced Materials 36, 2408770 (2024).

19. Hardin, J. O., Ober, T. J., Valentine, A. D. & Lewis, J. A. Microfluidic Printheads for Multimaterial 3D Printing of Viscoelastic Inks. Advanced Materials 27, 3279–3284 (2015).

20. Hassan, I. & Selvaganapathy, P. R. Microfluidic Printheads for Highly Switchable Multimaterial 3D Printing of Soft Materials. Adv Materials Technologies 7, 2101709 (2022).

21. Skylar-Scott, M. A., Mueller, J., Visser, C. W. & Lewis, J. A. Voxelated soft matter via multimaterial multinozzle 3D printing. Nature 575, 330–335 (2019).

22. Gervais, T., El-Ali, J., Günther, A. & Jensen, K. F. Flow-induced deformation of shallow microfluidic channels. Lab Chip 6, 500 (2006).

23. Elgack, M. E. & Abdelgawad, M. Characterization of the Dynamic Flow Response in Microfluidic Devices. Small Methods 9, 2401773 (2025).

24. Wu, C., Owsley, K. & Di Carlo, D. Rapid Software-Based Design and Optical Transient Liquid Molding of Microparticles. Advanced Materials 27, 7970–7978 (2015).

25. Wu, C.-Y. et al. Shaped 3D microcarriers for adherent cell culture and analysis. Microsyst Nanoeng 4, 21 (2018).

26. Wu, C.-Y. et al. Monodisperse drops templated by 3D-structured microparticles. SCIENCE ADVANCES (2020).

27. Werner, H. et al. Stopping a Multilayered Co-Axial Flow in a 3D Printed Microchannel with Cascaded Nozzles. Adv Materials Technologies 10, 2500203 (2025).

28. Sahin, M. A., Shehzad, M. & Destgeer, G. Stopping Microfluidic Flow. Small 20, 2307956 (2024).

29. Bong, K. W. et al. Compressed-air flow control system. Lab Chip 11, 743–747 (2011).

30. Paulsen, K. S. & Chung, A. J. Non-spherical particle generation from 4D optofluidic fabrication. Lab Chip 16, 2987–2995 (2016).

31. Li, Z., Liu, C. & Sun, J. Hydraulic–electric analogy for design and operation of microfluidic systems. Lab Chip 23, 3311–3327 (2023).

32. Smith, K., Retallick, A., Melendrez, D., Vijayaraghavan, A. & Heil, M. Modeling Graphene–Polymer Heterostructure MEMS Membranes with the Föppl–von Kármán Equations. ACS Appl. Mater. Interfaces 15, 9853–9861 (2023).

33. Zou, Y. et al. 3D Hydrodynamic Flow Lithography. Adv Funct Materials e77590 (2026) doi:10.1002/adfm.77590.

34. Yasmin, M. & Gupta, M. Thermodynamical Study of Alcoholic Solutions of Poly (ethylene glycol) Diacrylate and Poly (ethylene glycol) Dimethacrylate. Int. J. Thermo 15, 111–117 (2012).

